# Effects of basalt amendment and mycorrhizal inoculation on soil chemical properties and maize growth

**DOI:** 10.1101/2025.11.03.686277

**Authors:** Lucilla Boito, Jet Rijnders, Laura Steinwidder, Patrick Frings, Arthur Vienne, Mirthe Maes, Erik Verbruggen, Sara Vicca

## Abstract

Enhanced weathering (EW) of silicate minerals has emerged as a promising carbon dioxide removal (CDR) strategy, with potential benefits for soil fertility and crop performance. However, the soil processes that determine these co-benefits remain poorly constrained. In particular, interactions between basalt amendments and soil biota such as arbuscular mycorrhizal fungi (AMF) may influence nutrient mobilization and plant uptake, but these effects have rarely been quantified. In a 113-day mesocosm experiment with *Zea mays* using a Belgian, sandy loam soil, we investigated the effect of basalt and AMF inoculation on soil properties, nutrient and heavy metal availability, and crop yield and quality. We also assessed potential AMF-driven bio-weathering via cation mass balance and pore water dissolved inorganic carbon (DIC), pH, and alkalinity measurements. Basalt application, but not AMF, improved soil pH, cation exchange capacity, base saturation, and generally increased exchangeable Ca and Mg, whereas most other nutrients in the pore water remained unaffected. Crop yield and quality were largely unaltered by basalt or AMF, except for an increase in plant Mg with basalt application. Moreover, heavy metal availability and plant uptake were also generally unaffected, with the notable exception of soil pore water and corn Ni, which increased with basalt. These results suggest that risk for heavy metal contamination is not generic but may arise under specific environmental conditions. Finally, despite a synergistic effect of basalt and AMF on pore water DIC, we found no indication that AMF enhanced basalt weathering rates. Overall, AMF had limited influence on soil fertility indicators and crop performance. Basalt application improved key soil chemical indicators and increased the exchangeable fractions of Ca and Mg, demonstrating its role as a soil improver. Unlike several studies conducted in more acidic soils, these chemical enhancements did not increase maize growth here, indicating that the agronomic benefits of basalt are context-dependent.

## 1 Introduction

Amidst the urgent need for sustainable agricultural practices in the face of climate change (Branca et al., 2013), enhanced weathering (EW) of silicate minerals emerges as a promising solution, offering a novel approach to mitigate carbon dioxide (CO_2_) levels while simultaneously improving soil health and crop yields (Swoboda et al., 2022). Silicate minerals react with water and CO_2_ to form (bi)carbonate, resulting in long-term carbon (C) sequestration (Dietzen et al., 2018; Moosdorf et al., 2014). This is a naturally occurring process that regulates the global C cycle over geological timescales, but it is too slow to cope with the current rising atmospheric CO_2_ concentrations (Goudie and Viles, 2012). EW aims to accelerate this natural process by grinding the silicate rocks into a fine powder in order to increase their surface area and hence their weathering rate (Renforth et al., 2011). When applied to soils, this process is facilitated by moisture, elevated CO_2_ concentrations, and microbial activity.

Beyond its CDR potential, EW can deliver a suite of agricultural co-benefits. Silicate weathering releases nutrients, can counterbalance soil acidification, and improve soil water retention and cation exchange capacity (Swoboda et al., 2022; Van Straaten, 2006). Moreover, the cations liberated during weathering can facilitate the formation of stable mineral-associated organic matter (Slessarev et al., 2021), potentially improving soil fertility (Oldfield et al., 2019; Woomer et al., 1994). As a result, EW is generally expected to stimulate plant growth and crop yield (Goll et al., 2021; Shamshuddin et al., 2011; Vaghetti Luchese et al., 2021). However, the mechanisms governing these co-benefits remain insufficiently constrained, particularly the biotic interactions that may modulate EW effects in soils.

Basalt is a promising candidate for agricultural soil amendment due to its widespread availability and mineral composition. As an igneous rock rich in magnesium and iron silicates, it can also contain essential macronutrients such as calcium, phosphorus, and potassium, along with important micronutrients (Conceição et al., 2022). Compared to other (ultra)mafic silicate rocks like dunite, basalt contains lower concentrations of toxic heavy metals such as nickel and chromium, making it a safer option for agricultural use (Amann et al., 2020; Swoboda et al., 2022). The bioavailability of heavy metals is also strongly influenced by pH, with higher pH generally reducing the solubility and uptake of metals like nickel and zinc (Basta et al., 1993). Basalt application may thus improve soil fertility while minimizing contamination risks, although local soil conditions can impact these outcomes.

Among soil biota, arbuscular mycorrhizal fungi (AMF) play a central role in plant nutrition and soil processes. Approximately 80% of terrestrial plant species, including many important crops such as maize, wheat, and soybean, engage in a symbiotic relationship via their roots with arbuscular mycorrhizal fungi (AMF) (Verbruggen and Toby Kiers, 2010; Wilkes, 2021). AMF grow a soil hyphal mycelium that is specialized in nutrient and water uptake, and can reach a density of over 100 m of hyphae per cm³ of soil. These nutrients can be delivered to the host plant via arbuscules, AMF structures that are formed inside the root cells of the host plant (Parniske, 2008). In exchange, AMF receive carbohydrates that can be used for their growth (Kalamulla et al., 2022; Parniske, 2008; Verbruggen et al., 2021). Besides nutrient exchange, AMF can also influence soil structure and aggregation by exudation, turnover and enmeshment of soil particles (Rillig et al., 2015).

AMF can also affect the natural weathering of mineral rocks through several physical and chemical processes. Hyphae exert physical pressure on minerals, widening existing cracks and increasing the reactive surface area (Koele et al., 2014; Smits and Wallander, 2017). AMF also act as a sink for weathering products, and thus create chemical disequilibria across the rock surface which can enhance its weathering (Quirk et al., 2015; Verbruggen et al., 2021). During nutrient uptake, AMF release protons that acidify the soil surroundings, thus aiding the natural weathering process (Neumann and George, 2010; Taylor et al., 2009). AMF respiration also increases CO_2_ concentration in the soil (Moyano et al., 2007; Taylor et al., 2009), thereby creating a more favourable environment for rock weathering (Winnick and Maher, 2018). Moreover, AMF may influence the bioavailability and uptake of heavy metals through changes in soil chemistry (Riaz et al., 2021).

Despite the important role of AMF in promoting nutrient uptake as well as mineral weathering, experimental evidence on the interaction between AMF and EW is limited (Verbruggen et al., 2021). In this study, a mesocosm experiment was established with the aim of exploring the interaction between AMF and EW in agriculture. We hypothesized that i) AMF increases basalt weathering rates, and thus higher dissolved organic carbon (DIC) and alkalinity concentrations in the pore water and base cations in the soil-water complex are expected; ii) basalt application improves soil fertility and therefore plant growth and nutrient uptake, and that this effect is enhanced by AMF; iii) basalt addition does not increase heavy metal availability due to pH buffering, while AMF may modulate these effects by altering nutrient and metal transport between soil and plant; iv) AMF growth is affected by basalt application as a result of changes in soil nutrients and fertility.

## 2 Materials and Methods

### 2.1 Experimental set-up

Twenty mesocosms (0.6 m height, 0.25 m radius) were constructed at the experimental site at the Drie Eiken Campus of the University of Antwerp, Belgium (51⁰ 09’ N, 04⁰ 24’ E), and placed outdoors to receive natural rainfall. To allow leachate collection, a 2 cm hole at the bottom of each mesocosm was connected via polyurethane tubing to a glass collector (2.3 L). To prevent soil from leaching into the glass collector, a soil exclusion mat covered the bottom of each mesocosm.

In order to compare treatments with and without AMF, all soil was pasteurized by heating for 4 h at 80 °C (Ven et al., 2019; Verlinden et al., 2018). AMF colonization and hyphal length results at the end of the experiment confirmed that pasteurization succeeded, as treatments that were not inoculated (see further) showed clearly lower hyphal length and colonization (Fig. 1). The bottom 40 cm of all mesocosms was filled with a pasteurized sandy loam soil (69.5% sand, 28.1% silt, 1.8% clay) originating from a pasture in Zandhoven, Belgium. The initial soil was slightly acidic (pH = 5.5 ± 0.3), with an organic carbon content of 0.8 ± 0.05 % and a cation exchange capacity (CEC) of 3.9 ± 0.6 meq 100 g^-1^ soil. On 23 May 2022, the top 20 cm was filled with non-amended, pasteurized soil for the control treatments (n = 10 mesocosms). The top 20 cm of the basalt treatment (n = 10 mesocosms) received pasteurized soil with a mixture of 0.98 kg (equivalent to 50 ton ha^-1^) of freshly mined Eifel Basalt (Table 1; Rheinischen Provinzial-Basalt-un Lavawerke (RPBL)), with particle size between 0.02 and 0.2 mm (Fig. S1) and specific surface area (SSA) corresponding to 6.283 ± 0.001 m^2^ g^-1^ (mean ± SE) measured on three technical replicates with a Quantachrome Autosorb iQ using the Braunauer-Emmet-Teller method. The measurement used nitrogen (N_2_) as adsorbate with multi-point (5 points) and isotherm (77K) settings. The samples were degassed at 300°C with 200 minutes of soak time. High measurement quality was ensured by frequent reference (Bundesanstalt für Materialforschung und –prüfung) measurements.

**Figure 1:**
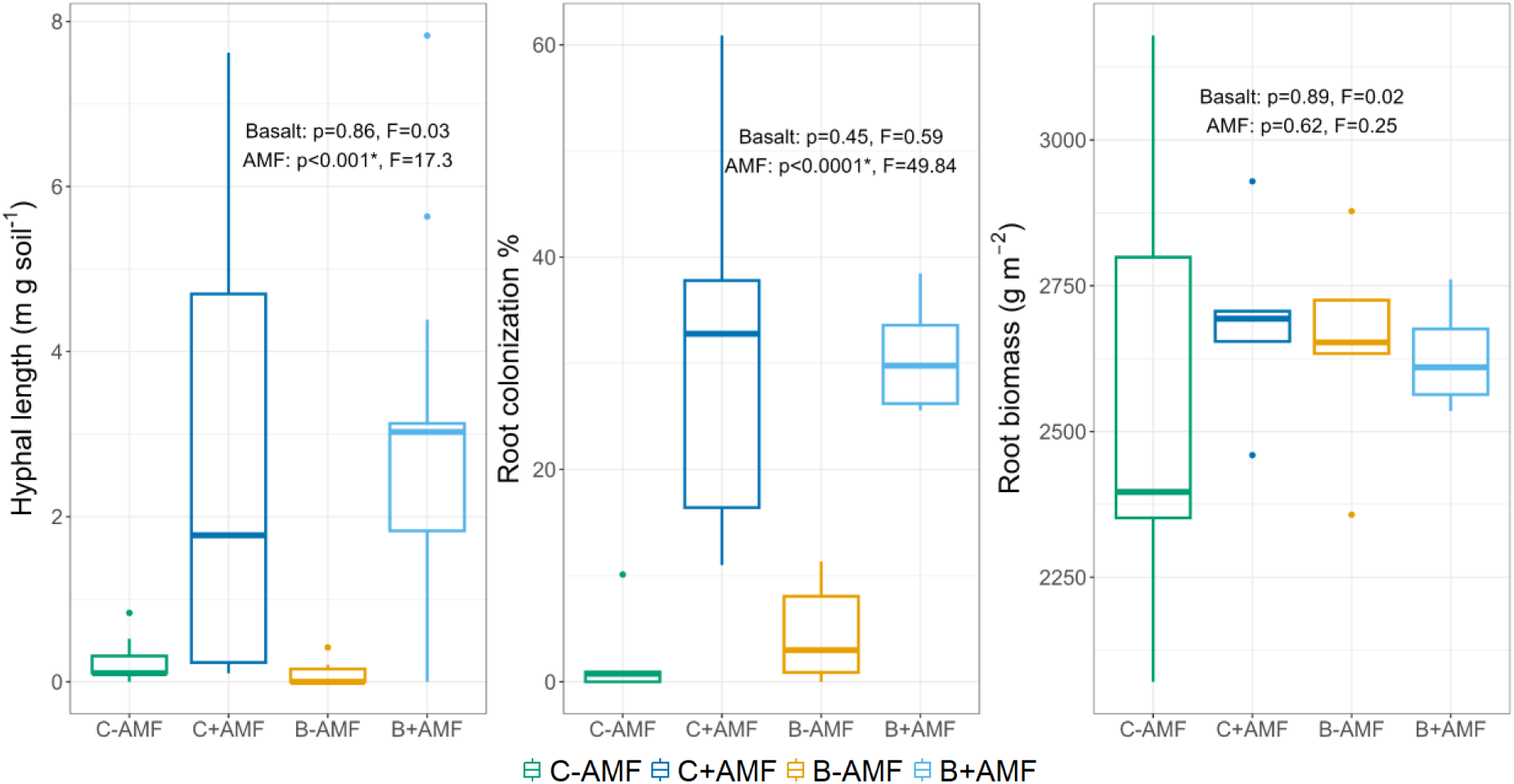
AMF hyphal length, root colonization by AMF and root biomass of the four treatments (C-AMF = control without AMF, C+AMF = control with AMF, B-AMF = basalt without AMF, B+AMF = basalt with AMF) at the end of the experiment. Boxes represent the interquartile range (25th– 75th percentile), horizontal lines indicate the median, and whiskers extend to the most extreme data points within 1.5× of the interquartile range. Points outside this range are plotted as outliers. p– and F-values from a linear regression analysis with hyphal length, root colonization or biomass as response variables and basalt, AMF and their interaction as covariables are shown. Interactions were not significant and were excluded from the model. Statistical significances are indicated with an asterisk (*).

**Table 1:** Basalt mineral phases obtained via XRD analysis and elemental concentrations obtained via XRF analysis. C = concentration.

| Mineral phase | Mass % |  | Element | C (wt%) | Element | C (wt%) |
| --- | --- | --- | --- | --- | --- | --- |
| Clinopyroxene | 21.22 vs |  | Si | 41.95% | Ni | 394 PPM |
| Enstatite | 0.66 |  | Mg | 14.58% | Cr | 442 PPM |
| Pigeonite | 2.36 |  | Fe | 12.87% | V | 211 PPM |
| Forsterite | 14.45 |  | Al | 10.93% | Zr | 201 PPM |
| Hedenbergite | 6.98 |  | Ca | 10.05% | Zn | 120 PPM |
| Ankerite | 0.18 |  | Ti | 2.11% | Ce | 103 PPM |
| Grossmanite | 1.29 |  | Na | 1.73% | Nb | 68.7 PPM |
| Albite | 1.29 |  | K | 0.91% | Co | 60.3 PPM |
| Labradorite | 5.7 vs<br>7.49 |  | P | 0.61% | Cu | 54.3 PPM |
| Analcime | 0.93 |  | Mn | 0.22% | Rb | 28.0 PPM |
| Quartz | 0.07 |  | S | 0.04% | Y | 25.3 PPM |
| Magnetite | 2.73 |  | Cl | 0.03% |  |  |
| Amorphous phase | 42.14 |  | Ba | 525 PPM |  |  |

On 15 June 2022, all mesocosms were planted with two dwarf corn plants each (*Zea mays* L., *golden midget* variety) to mimic a field density of about 13 plants m^-2^. For half of the treatments (five control and five basalt mesocosms), plants were inoculated with spores of arbuscular mycorrhizal fungi (AMF; species *Rhizophagus irregularis*; Symplanta), while the remaining 10 mesocosms (five control and five basalt treatments) received a pasteurized inoculum (Table 2). In addition, a microbial wash was added to every plant to avoid initial differences in microbial community due to inoculation (Wagg et al., 2011). The microbial wash was obtained by suspending 20 g of dry inoculum in 200 ml of deionized water. After mixing, the liquid inoculum was filtered through a Whatman Grade 41 ashless filter paper (12.5 cm diameter, nominal retention: 20–25 µm, Cat. No. 1441 125, Cytiva), and a few drops were inspected under a microscope to ensure the absence of AMF spores. All pots were fertilized with nitrogen, phosphorous and potassium (NPK; 96 – 10 – 39.5 kg ha^-1^) by adding ammonium nitrate (NH_4_NO_3_), triple super phosphate (TSP; 45% P_2_O_5_) and potassium sulphate (K_2_SO_4_). The N fertilization amount was similar to that used in Ven et al. (2019), whereas P and K application amounts were halved to avoid overfertilization in combination with the added silicates.

**Table 2:** Treatments of the experiment with number of replicates and abbreviations.

| Treatment | Meaning | # replicates |
| --- | --- | --- |
| C-AMF | Control treatment without AMF | 5 |
| C+AMF | Control treatment with AMF | 5 |
| B-AMF | 50 ton ha<sup>-1</sup> of basalt without AMF | 5 |
| B+AMF | 50 ton ha<sup>-1</sup> of basalt with AMF | 5 |

To avoid excessive heating, mesocosms were wrapped in reflective foil. One mesh bag (13 cm x 3 cm x 2 cm) with a mesh size of 30 µm, which is permeable to AMF hyphae but excludes roots, was buried in each mesocosm at a depth of 0-10 cm. The mesh bags were filled with white sand that was preheated in the oven at 550 °C for 5h to remove organic matter.

During the experiment, several plants were infected with corn smut, a disease caused by the pathogenic fungus *Ustilago maydis*. The fungus forms tumours on the aerial parts of its host plant (Matei and Doehlemann, 2016) and can significantly decrease maize yield (Clough and Blatchford, 2011). Whenever a plant was infected and a tumour was observed, the tumour was removed from the plant. Fresh weight was determined, whereafter the tumour was dried in the oven at 70 °C for 48h and dry weight was measured. Corn smut weight did not differ among the treatments (Table S2).

### 2.2 Soil fertility and pore water measurements

From 1 June 2022, soil pore water samples were collected every two weeks using rhizon samplers (Rhizon Flex, Rhizosphere Research Products B.V.) installed vertically at 0-10 cm depth in each mesocosm. Pore water analyses included alkalinity, pH, and concentrations of dissolved inorganic carbon (DIC), phosphate (PO_4_ ^3-^), and the metals aluminium (Al), calcium (Ca), iron (Fe), potassium (K), magnesium (Mg), sodium (Na), nickel (Ni), silicon (Si) and zinc (Zn). Pore water ammonium (NH_4_ ^+^-N), nitrate (NO_3_ ^-^ –N), sulphate (SO_4_ ^2-^), copper (Cu) and manganese (Mn) concentrations were also quantified and are reported in Figure S3. Soil pore water pH was measured with a 914 pH/Conductometer (2.914.0020, Metrohm). DIC was measured on three technical replicates with a TOC analyser (Formacs^TM^ HT Series, Skalar) using deionized water as blank and sodium carbonate (EMSURE, Supelco) as quality control; whereas alkalinity, nitrogen, SO_4_ ^2-^ and PO ^-^ were measured on three technical replicates by a continuous flow analyser (SAN++® Advanced Series, Skalar) using sodium carbonate (Certipur, Merck Group) and standard NH_4_ ^+^, NO_3_ ^-^, PO_4_ ^3-^ and SO_4_ ^2-^ solutions (Certipur, Supelco) as quality controls respectively. DIC, nitrogen, PO_4_ ^3-^ and SO_4_ ^2-^ analyses were conducted in-house on samples filtered through a disposable 0.45 µm PET-45/25 membrane filter (CHROMAFIL, Macherey-Nagel). Elemental analyses were performed by ICP-OES (DSOI, SpectroGreen, SPECTRO) in the Helmholtz Laboratory for the Geochemistry of the Earth Surface (HELGES) on samples filtered through a disposable 0.2 µm PET-20/25 membrane filter (CHROMAFIL, Macherey-Nagel) using international water reference material SLRS-6 (NRC-CNRC) as quality control. Alkalinity measurements were carried out in-house on unfiltered samples.

Measurements of soil fertility (here defined as soil pH, CEC and base saturation) were performed on topsoil samples collected before basalt application and at the end of the experiment (12 September 2022). Topsoil pH was measured by collecting, pooling and air-drying three subsamples per mesocosm. Subsequently, 4 g of soil was dissolved in 10 mL deionized water and shaken before measuring using a 914 pH/Conductometer (2.914.0020; Metrohm). Soil CEC was determined after Brown (1943).

### 2.3 Sequential extractions of soil cations

Sequential extraction techniques are designed to quantify elements in different soil fractions (Tessier et al., 1979). These are valuable for calculating weathering rates of the silicate minerals (Niron et al., 2024; Vienne et al., 2024) as well as to quantify cations available for plant uptake (i.e. in the soil exchangeable complex). For this, soil samples were collected at the beginning of the experiment (day 9 after basalt amendment) at a depth of 0-20 cm (sampling deeper soil layers was not possible as it would entail destructive sampling). At the end of the experiment (day 113), soil samples were taken at three depth intervals (thus defining three soil layers: 0-20 cm, 20-40 cm, and 40-60 cm) and subsequently air-dried.

Sequential extractions were performed using protocols adapted from Tessier et al. (1979) and Uhlig and Von Blanckenburg (2019). In a four-step process, Al, Ca, Cr, Fe, K, Mg, Na, Ni, Si, and Zn were extracted from four soil pools, i.e., the exchangeable, carbonate, reducible and oxidizable pool. Cations that remained in the soil after the extraction process are considered to be part of the primary or secondary minerals’ structure, and thus unlikely to be released under typical soil conditions (Tessier et al., 1979). The concentrations of Al, Ca, Cr, Fe, K, Mg, Na, Ni, Si, and Zn for each extraction step were then measured in the Helmholtz Laboratory for the Geochemistry of the Earth Surface (HELGES) on three technical replicates by ICP-OES (DSOI Spectrogreen, SPECTRO), using international water reference material SLRS-6 (NRC-CNRC) as quality control. In order to exclude the cations extracted from the initial soil and un-weathered basalt, and investigate the effect of basalt and AMF on the cations in the various soil fractions, soil cation concentrations at the start of the experiment were subtracted from the concentrations measured at the end of the experiment. In the main text only cation concentrations in the exchangeable fraction are reported and widely discussed. These are cations adsorbed onto the soil’s exchangeable complex, and are therefore relevant for soil fertility, plant uptake and growth. Results from reducible and oxidizable pools are reported in the appendix (Fig. S4-5; Table S6-7) and are only briefly discussed in the main text, as they provide indication on clay-sized mineral and stable organic C formation respectively.

### 2.4 Weathering rates calculations

Weathering rates were calculated based on the major cations (Ca, Mg, Na, K) released from the applied basalt (as in Steinwidder et al., 2025). These cations can follow several paths: they either leach out of the soil, are taken up by plants, or remain within the soil system. During the experiment, high temperatures were registered, and leaching occurred only at the beginning of the experiment and in minimal volumes (Table S8), as most of the water in the mesocosms was lost through evapotranspiration driven by plant uptake and atmospheric evaporative demand. The leaching loss of cations was thus negligible and only the cations accumulating in plant biomass and soil pools were used for the weathering rate calculations.

To determine the amount of each cation released from the applied basalt and taken up by plants, cation concentrations in the C-AMF treatment were subtracted from cation concentrations in basalt-applied treatments. Average leaf cation concentration was used to calculate the increase in mol charge equivalents with basalt application compared to C-AMF treatment.

In order to determine the amount of each cation released into the soil from the weathering of the applied basalt over the experimental duration, initial (day 9) cation concentrations in each sequentially extracted pool were subtracted from the final cation concentrations extracted at the experiment’s conclusion (day 113). As the extraction process also extracts primary minerals from un-weathered basalt, this approach accounts for the contribution of the un-weathered basalt to the extracted cations pool by subtracting the initial (mostly un-weathered) soil + basalt mixture from the final concentrations. Finally, in order to account for potential changes over time in the soil due to plant uptake and/or inclusion of cations within clays, the increases in cation concentration in the basalt treatments were expressed relative to the C-AMF treatment.

For the 0-20 cm layer, initial cation concentrations were subtracted from the final concentrations. For the deeper layers, where no basalt was incorporated, initial cation concentrations were assumed to be equivalent to initial concentrations in the top layer of the control treatment without basalt (C-AMF). Any cation increase observed in the lower layers was therefore attributed to weathering products, as un-weathered basalt was absent in these layers. This calculation assumes that cation migration to the deeper soil layers occurs exclusively as a result of weathering. Weathering rates (Wr; expressed as log_10_ (mol charge equivalents m^-2^ s^-1^)) were then calculated as in Steinwidder et al. (2025) (Eq. 1):

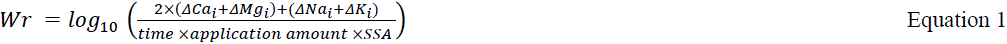

Where ΔCa_i_, ΔMg_i_, ΔNa_i_, and ΔK_i_ are the concentration changes in each soil fraction (i.e. exchangeable, carbonate, reducible and oxidizable pools) and plant pool *i* calculated as mentioned above, time is the experimental duration in seconds, application amount is 980 g mesocosm^-1^ (equivalent to 50 ton ha^-1^) and SSA is the specific surface area of the applied basalt, with a value of 6.283 m^2^ g^-1^ (see section 2.1).

### 2.5 Plant measurements

Plants were harvested on 12 September 2022. The three top leaves of each plant were sampled for further analysis, whereafter the aboveground biomass was harvested and separated into stems, leaves, tassels and corn. Aboveground biomass was then dried at 70 °C for 48 h and the dry weight of each organ was determined. After harvesting the aboveground biomass, roots were sampled from the topsoil (0-20 cm), middle soil layer (20-40 cm) and bottom soil (40-60 cm) to estimate the root biomass within each mesocosm. For each layer, two 100-cm^3^ cores were taken directly beneath each plant and one between the two plants. The roots were then washed using a 1 mm mesh sieve, after which they were dried at 70 °C for 48h and weighed. Root biomass estimates assumed that cores beneath the plants represented root growth in half of the mesocosm, while cores between the plants reflected the other half (as in Ven et al., 2020). Before analysis, the three top leaves (hereafter referred to as leaves), stems, tassels, corn kernels and roots were ground using a centrifugal mill (ZM 200, Retsch GmbH) with a 0.25 mm mesh size sieve.

The ground samples were analysed for nutrients (C, Ca, Fe, K, N, Mg, P) and heavy metals (Al, Cd, Cr, Ni, Pb, V and Zn) by ICP-OES (iCAP 6300 Duo, Thermo Scientific) using certified standards (CPAChem) as quality controls every 20 samples and Hay Powder BCR129 as Reference Material (CPAChem). Plant Ni, Cr and Mg results are reported in the main text, while concentrations of all other elements are reported in the supplements (Fig. S9). Biogenic Si was determined by digestion of a 0.03 g plant sample with 25 mL O.5 N NaOH and then analysed with a continuous flow analyser (SAN++® Advanced Series, Skalar) following Demaster (1981); and Saccone et al. (2007).

### 2.6 AMF measurements

To determine plant root colonization by AMF, plant roots were first cleared and then stained using a non-vital staining technique (Vierheilig et al., 2005). Washed roots were cleared by placing them in an Erlenmeyer flask containing a 5% KOH solution which was then set in a warm water bath at 90 °C for 8 minutes. Hereafter, the roots were rinsed and stained with 10% Sheaffer Black ink in 10% acetic acid by placing them in a warm water bath at 90 °C for 10 minutes. To remove excess ink, the roots were immersed in lactoglycerol (1:1:1 lactic acid:glycerol:deionized water) and stored in an Eppendorf tube with lactoglycerol overnight at 4 °C. The proportion of roots colonized with mycorrhizal fungi was then quantified by counting arbuscules, vesicles and hyphae using the gridline intersection method (Fig. S10-11) (Giovannetti and Mosse, 1980).

During the harvest on 12 September 2022, the mesh bags were removed from the mesocosms and stored frozen until further analysis (Fig. S11a). Hyphal length of the mycorrhizae was subsequently determined according to Rillig and Field (1999) with adaptations as in Ven et al. (2019).

### 2.7 Data analysis

All statistical analyses were performed in R using RStudio packages nlme, lmerTest, dunn.test (Kuznetsova et al., 2017; Pinheiro et al., 2007; R Core Team, 2024; v4.3.1; Dinno, 2024). The level of significance for all analyses was set at p < 0.05. A factorial analysis of variance (ANOVA) was used to investigate the effects of basalt application, AMF inoculation, and their interactions, on soil pH, CEC, base saturation, exchangeable cation concentration in each soil layer and weathering rates. If normality or homoscedasticity of the model residuals were not met after data transformation, the non-parametric Kruskal-Wallis test was used with treatment (C-AMF, C+AMF, B-AMF and B+AMF) as explanatory variable. A post-hoc test (Tukey’s honestly significant difference test or Dunn’s multiple comparison test for parametric and non-parametric analyses respectively) was conducted to explore significant differences among treatments.

When cation concentrations in soil were below the detection limit (=bd; Table S12-13), the concentrations were set to zero to allow further statistical analyses. In some cases, the presence of the cation was detectable, but the very low concentration did not allow an accurate quantification. For these measurements, half of the limit of quantification (LOQ/2) was used for statistical analyses.

To quantify the treatment effect on variables with repeated measurements over time, i.e. pore water pH, DIC, alkalinity, and nutrient and heavy metal concentrations in the pore water, a linear mixed model was employed. Basalt application, AMF, and time (days after basalt application), and their interactions were included as fixed effects, and mesocosm was included as a random effect term. To assess whether the presence of galls from corn smut was affected by basalt, AMF or their interaction, a generalized linear model was used with the binomial family, as appropriate when modelling binary response variables.

The data and model code in support of our findings are openly available in Zenodo at https://doi.org/10.5281/zenodo.16813184 (Boito et al., 2025).

## 3 Results

### 3.1 Basalt improved soil fertility, but did not affect most pore water nutrients

At the end of the experiment, basalt significantly increased soil pH, CEC, and base saturation (Fig. 2). The influence of basalt on soil pH, CEC, and base saturation was not affected by AMF. Moreover, pore water pH and alkalinity also increased with basalt, without influence of AMF (Fig. 3). Concerning DIC, our data showed significant two– and three-way interactions between basalt and time and between basalt, AMF, and time respectively. Basalt increased DIC concentrations in the pore water over time, and this effect was reinforced by AMF towards the end of the growing season (Fig. 3).

**Figure 2:**
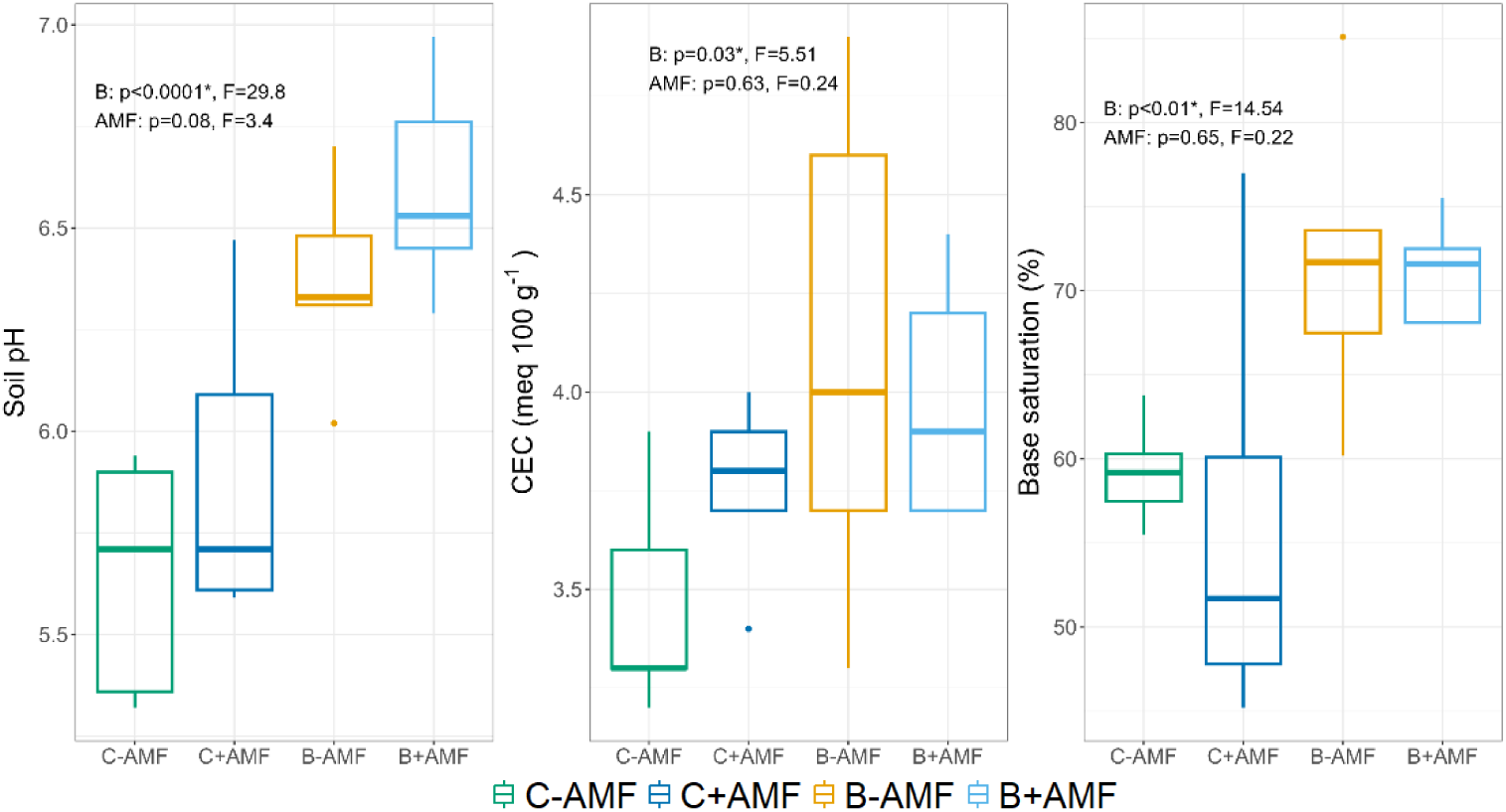
Soil pH, cation exchange capacity (CEC) and base saturation of the four treatments. (C-AMF = control without AMF, C+AMF = control with AMF, B-AMF = basalt without AMF, B+AMF = basalt with AMF) at the end of the experiment. Boxes represent the interquartile range (25th–75th percentile), horizontal lines indicate the median, and whiskers extend to the most extreme data points within 1.5× of the interquartile range. Points outside this range are plotted as outliers. p– and F-values from a linear regression analysis with pH, CEC or base saturation as response variables and basalt, AMF and their interaction as covariables are shown. Interactions were not significant and were excluded from the model. Statistical significances are indicated with an asterisk (*).

**Figure 3:**
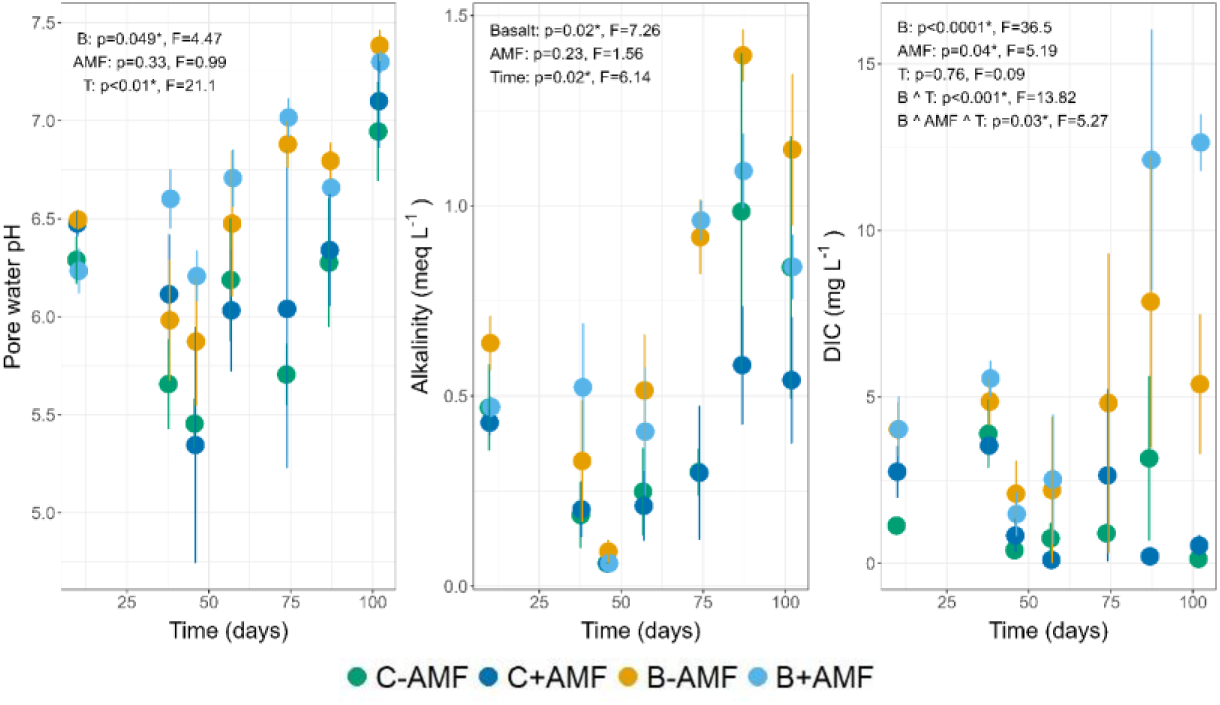
Mean pH, total alkalinity and dissolved inorganic carbon (DIC) in the soil pore water during the experiment for the four treatments (C-AMF = control without AMF, C+AMF = control with AMF, B-AMF = basalt without AMF, B+AMF = basalt with AMF). Error bars represent the standard error on the mean. p– and F-values from a linear regression analysis with pH, total alkalinity or DIC as response variables and basalt, AMF and time, and their relevant interactions, as covariables are shown. When interactions were not significant, they were excluded from the model and not shown here. Interactions are indicated by carets (^), while significant relationships by an asterisk (*).

Surprisingly, concentrations of plant-available nutrients in the pore water were generally not significantly affected by basalt application nor AMF presence, with the exception of basalt-driven increases in Mg and Na in the middle of the growing season (Fig. 4). Pore water Mg also showed a significant three-way interaction effect of basalt, AMF and time, with AMF reinforcing the positive effect of basalt (Fig. 4).

**Figure 4:**
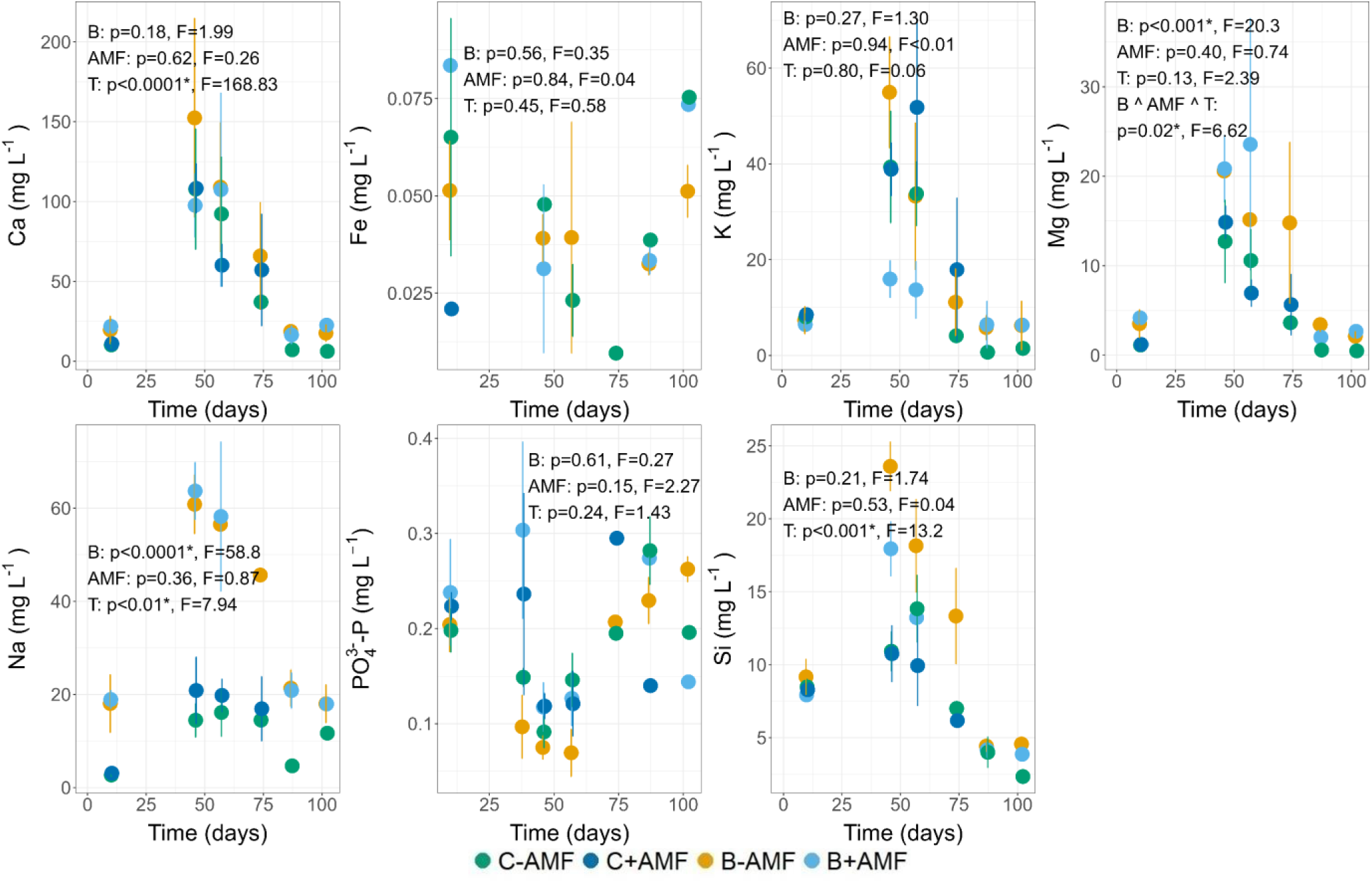
Mean concentrations of elements and nutrients in the pore water (Ca, Fe, K, Mg, Na, PO ^3-^ and Si) during the experiment for the four treatments (C-AMF = control without AMF, C+AMF = control with AMF, B-AMF = basalt without AMF, B+AMF = basalt with AMF). Error bars represent the standard error on the mean. p– and F-values are shown from a linear regression analysis with element concentration as response variable and basalt, AMF and time, and their relevant interactions, as covariables. When interactions were not significant, they were excluded from the model and not shown here. Interactions are indicated by carets (^), while significant relationships by an asterisk (*).

Concerning heavy metals, Cr in the pore water was below the limit of quantification (<0.02 mg L^-1^ for pore water concentrations) in all samples and for Ni, 67% of the samples were below LOQ, precluding formal statistical analysis. Nonetheless, basalt tended to increase Ni concentration, albeit only in the first half of the growing season (Fig. 5). A significant time x basalt interaction was found on Zn concentrations in the pore water, although without a clear trend. Pore water Al decreased with basalt amendment, and AMF presence decreased pore water Al but only in control mesocosms (significant basalt x AMF interaction).

**Figure 5:**
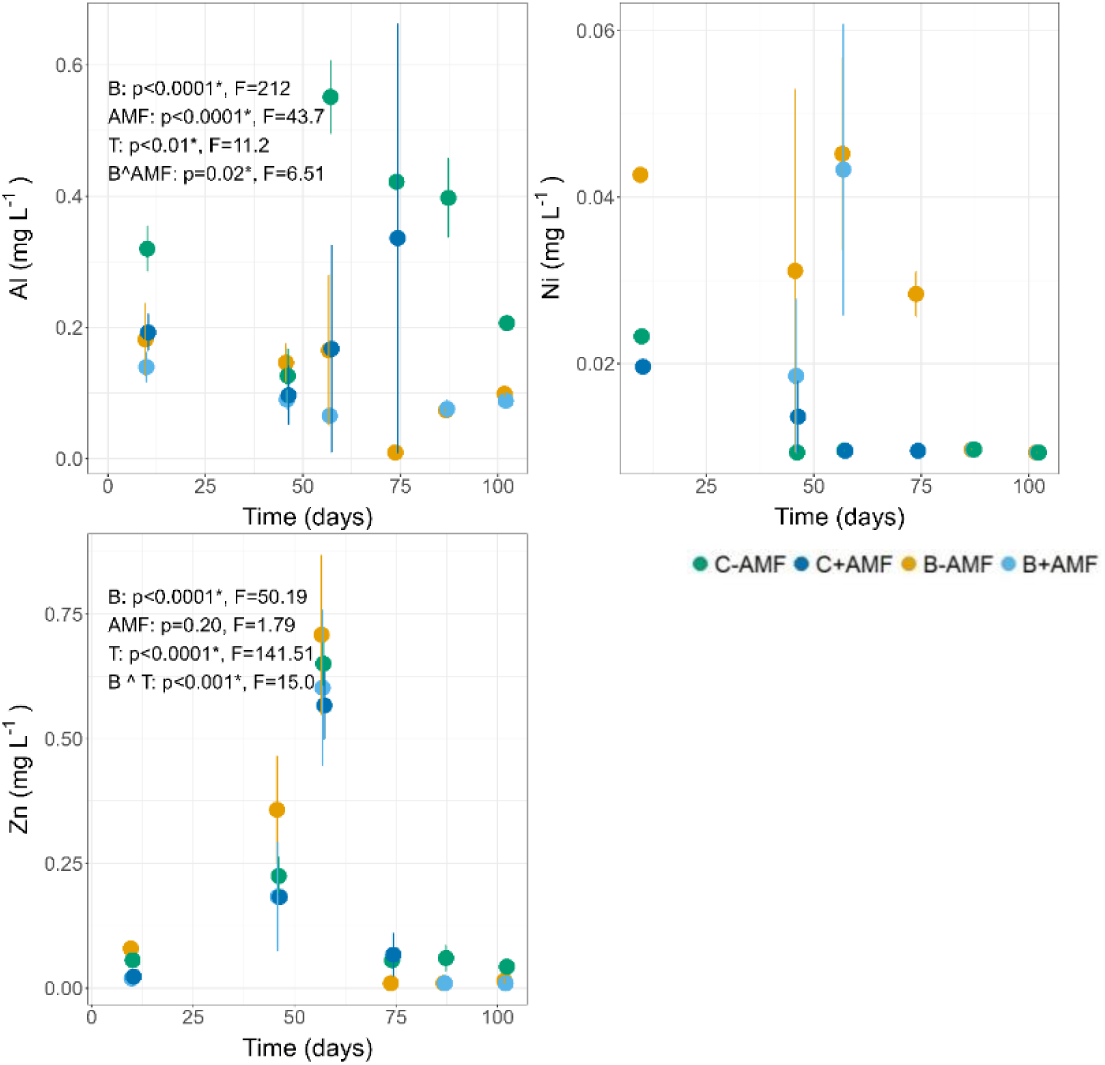
Pore water heavy metal concentrations. (Al, Ni, and Zn) during the experiment for the four treatments (C-AMF = control without AMF, C+AMF = control with AMF, B-AMF = basalt without AMF, B+AMF = basalt with AMF). Error bars represent the standard error on the mean. p– and F-values are shown from a linear regression analysis with heavy metal concentration as response variable, and basalt, AMF and time, and their relevant interactions, as covariables. When interactions were not statistically significant, they were excluded from the model and not shown here. Interactions are indicated by carets (^), while significant relationships by an asterisk (*).

#### 3.1.1 Basalt increased exchangeable Ca, Mg and Na, while it decreased exchangeable Zn

Basalt application increased exchangeable Ca and Mg in the 0-20 and 20-40 cm layers but not in the 40-60 cm layer (Fig. 6; Table S6). Additionally, basalt increased exchangeable Na in the 20-40 cm layer, and in the 0-20 cm layer a post-hoc pairwise comparison revealed a significant Na increase between C-AMF and B+AMF treatments. On the other hand, exchangeable K, Fe and Si concentrations were not affected by basalt application (Fig. 6; Table S6). Concentrations of exchangeable Ni and Cr were below LOQ in all layers, and the same was true for Al in the 40-60 cm layer (Table S12), hence precluding formal statistical analysis of these elements in the exchangeable pool of those layers. This was not the case for exchangeable Al in the 0-20 and 20-40 cm layers, which was not affected by basalt application, and exchangeable Zn, which decreased in the 0-20 and 20-40 cm layers (Fig. 7; Table S6). AMF did not affect cation concentrations in the sequentially extracted soil pools.

**Figure 6:**
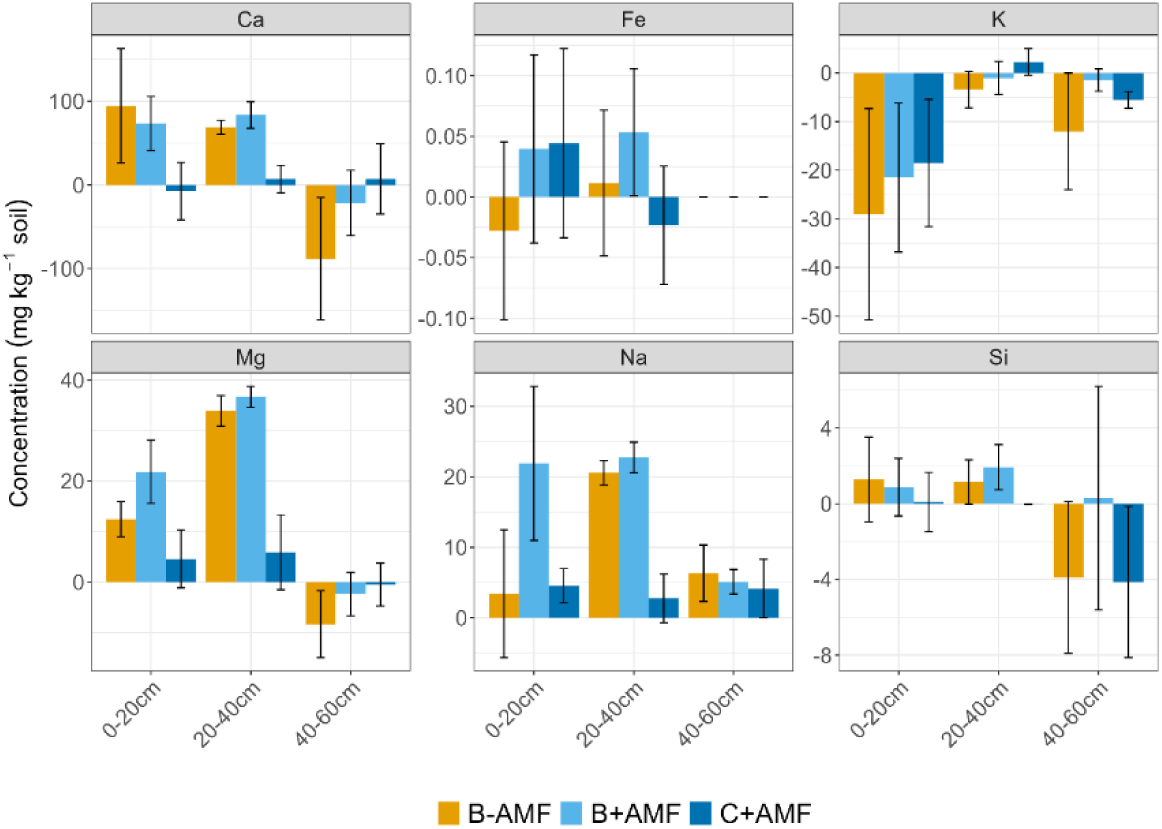
Mean soil nutrients (Ca, Fe, K, Mg, Na and Si) for the three treatments (C+AMF = control with AMF, B-AMF = basalt without AMF, B+AMF = basalt with AMF) at the end of the growing season. Error bars represent the standard error on the mean. Nutrient concentrations are shown as the difference between the start and the end of the growing season, and are shown compared to the control treatment (C-AMF; control without AMF). Soils were divided into three sampling depths (0-20 cm, 20-40 cm, 40-60 cm) for which nutrient concentrations were measured in the exchangeable fraction with sequential extraction method after Tessier et al. (1979). p– and F-values from a linear regression analysis with nutrient concentration as response variables, and basalt, AMF and their interaction as covariables are presented in table S6.

**Figure 7:**
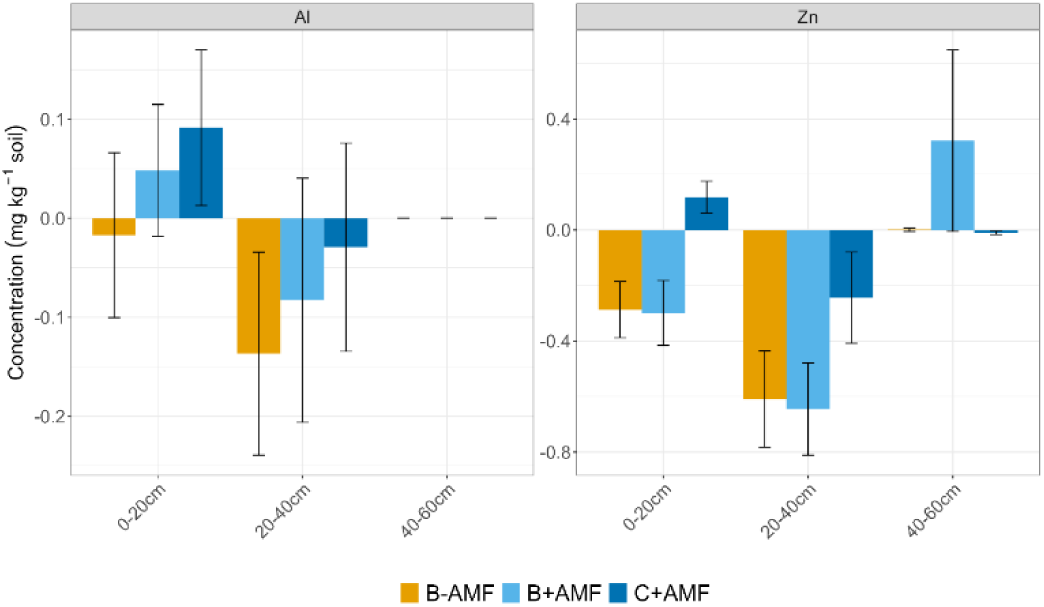
Mean soil heavy metal concentrations (Al and Zn) for the three treatments (B-AMF = basalt without AMF, B+AMF = basalt with AMF, C+AMF = control with AMF) at the end of the growing season. Error bars represent the standard error on the mean. Heavy metal concentrations are shown as the difference between the end of the growing season and the start, and are shown compared to the control treatment (C-AMF; control without AMF). Soils were divided into three sampling depths (0-20 cm, 20-40 cm, 40-60 cm) for which heavy metal concentration in the exchangeable fraction was measured with the sequential extraction method after Tessier et al (1979). p– and F-values from a linear regression analysis with nutrient concentration as response variables, and basalt, AMF and their interaction as covariables are presented in table S6.

### 3.2 AMF did not affect basalt weathering rates

Similarly, basalt weathering rates were not affected by AMF presence (Fig. 8), with rate constants corresponding to –9.97 ±0.044 and –10.01 ±0.053 log_10_(mol charge equivalents m^-2^ s^-1^) for treatments with and without AMF, respectively. This corresponds to the weathering of 37.5% ±3.7 and 33.2% ±4.0 of Ca-, Mg-, Na– and K-bearing minerals in the applied basalt, with and without AMF respectively, over the experimental duration (113 days). When comparing Mg, Ca, Na and K changes in soil pools with changes in plant pools, it emerged that the greatest contribution to the increase in released cations was in the soil phase, with a total increase in the soil about two orders of magnitude greater than in the plants (Table S14).

**Figure 8:**
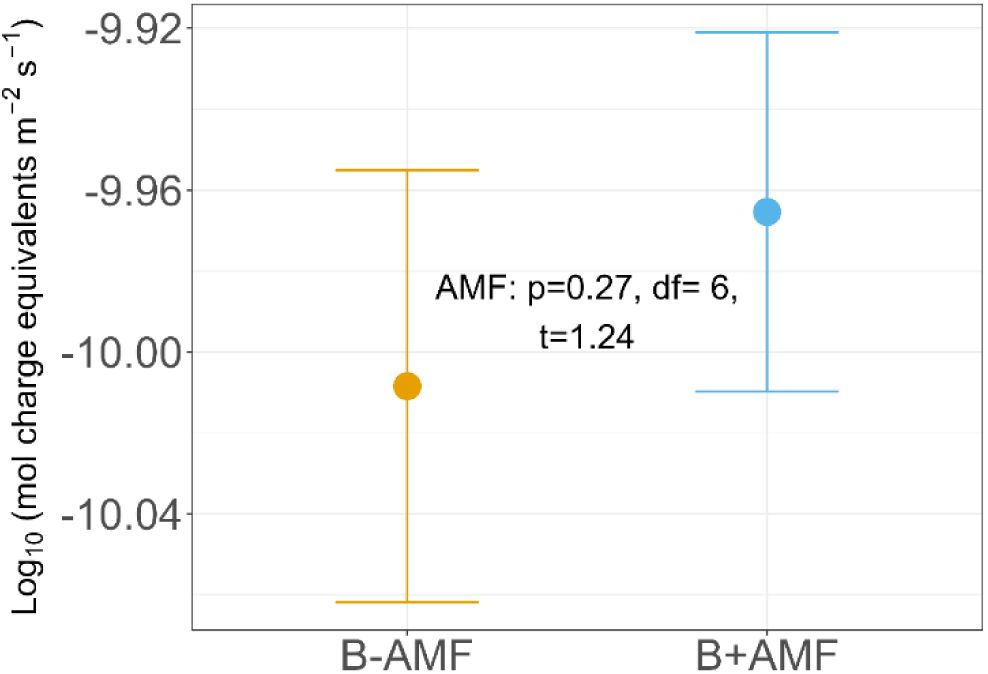
Mean basalt weathering rates for the basalt treatments calculated as the increase in base cations (Ca, Mg, K, Na) in the soil phase and in the plants between the start and the end of the experiment, and corrected for the control soil (C-AMF; control without AMF). Error bars represent the standard errors on the mean. p– and t-values from a two-sample t-test are presented.

### 3.3 Limited basalt and AMF effects on crop growth, biomass and quality

Generally, plant growth parameters were not affected by basalt or AMF, including measurements of individual plant organs (Fig. S15-16). In addition, no effect of basalt on AMF hyphal length and root colonization was found (Fig. 1). Corn smut presence did not differ among treatments, nor did the weight of the galls (Table S2). Elemental concentrations (Ca, Fe, Mg, K, Si, P) in plant parts were typically not affected by basalt or AMF application, with a few exceptions. A notable exception was Mg, whose concentration in plant organs was generally higher with basalt application, except for corn Mg where no basalt effect was found (Fig. 9; Table S17). Moreover, in contrast with the other elements in other plant organs, a significant basalt x AMF interaction effect was found on root Mg concentrations, which only increased with basalt when AMF were present. Moreover, basalt decreased leaf K, whereas AMF decreased leaf P. Finally, stem and tassel Si concentrations increased with basalt application (Fig. S9; Table S17).

**Figure 9:**
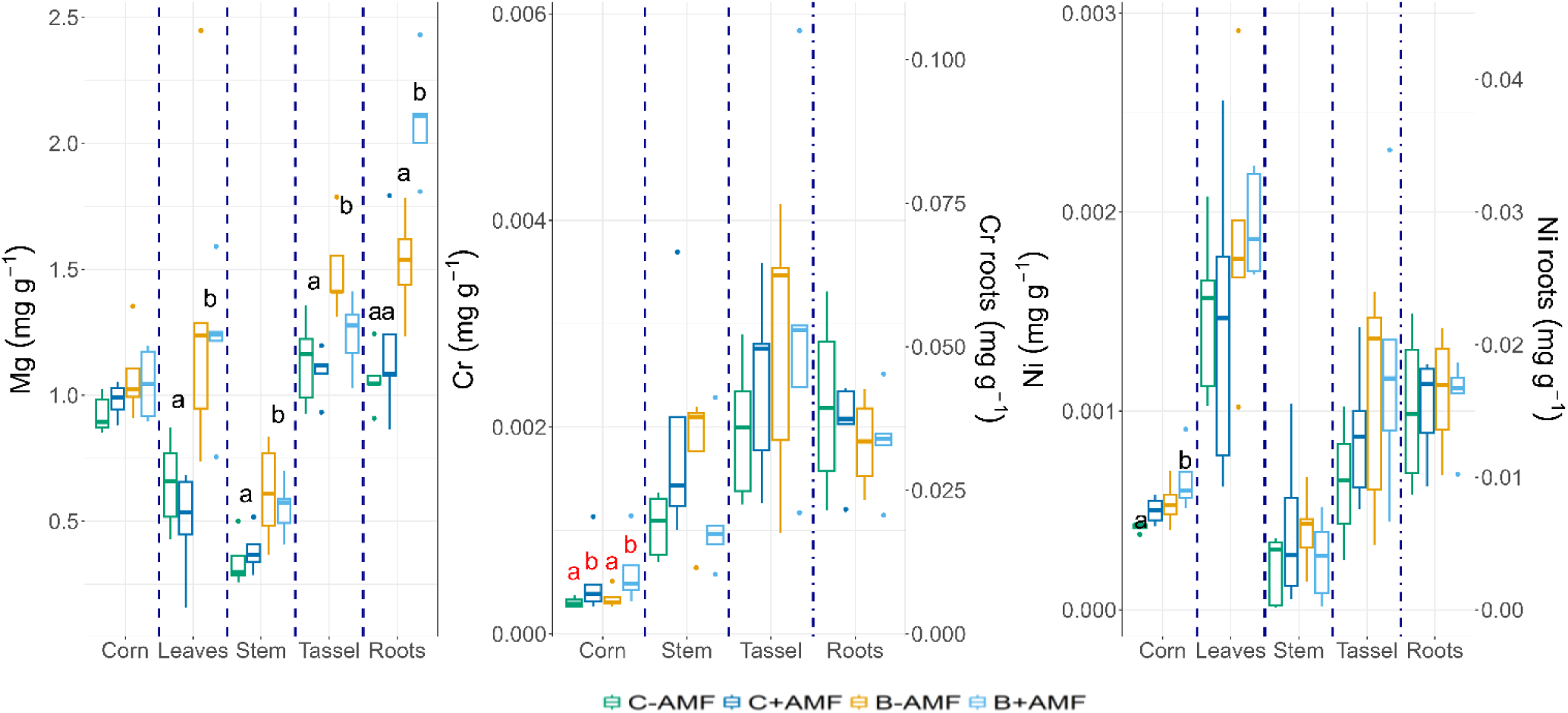
Concentrations of Mg, Cr and Ni in the corn, leaves, stem, tassel, and roots for the four treatments (B-AMF = basalt without AMF, B+AMF = basalt with AMF, C-AMF = control without AMF, C+AMF = control with AMF) at the end of the experiment. Boxes represent the interquartile range (25th–75th percentile), horizontal lines indicate the median, and whiskers extend to the most extreme data points within 1.5× of the interquartile range. Points outside this range are plotted as outliers. Note that for Cr and Ni, the primary axis shows the concentration in the corn, leaves, stems, and tassels while the secondary axis shows the concentration in the roots. p– and F-values are shown in table S17. Significant differences are indicated with a different letter, while a similar letter means no differences between those treatments. Black letters indicate basalt effects, while red letter indicate AMF effects. If no letters are shown for a combination of plant part and element, this means that there was no significant influence of basalt or AMF.

Leaf Cr concentrations were all below LOQ (<0.0002 mg g^-1^). Therefore, statistical analyses could not be conducted for leaf Cr. Basalt application did not affect Al, Cr and Zn concentrations in the various plant organs (Fig. 9; Fig. S9; Table S17). In contrast, corn Ni concentrations were significantly higher with basalt application (Fig. 9; Table S17). AMF also increased corn Ni concentrations significantly, but a post-hoc pairwise comparison revealed that only the C-AMF – B+AMF difference was significant. Finally, AMF increased corn Cr (Fig. 9; Table S17).

## 4 Discussion

### 4.1 Weathering rates and proxies

The weathering rates in our experiment (–9.97 ±0.044 and –10.01 ±0.053 log _10_ (mol charge equivalents m^-2^ s^-1^), with and without AMF respectively) are more than one order of magnitude higher than the weathering rates calculated based on cation tracing in similar experiments. For example, in a parallel trial at the same experimental location, Steinwidder et al. (2025) found weathering rates of –11.15 ±0.16 log_10_ (mol charge equivalents m^-2^ s^-1^), calculated with the same methodology, 15 months after the application of 50 ton ha^-1^ basalt on the same soil type, albeit unpasteurized. Another mesocosm experiment by Vienne et al. (2024) yielded a mean log weathering rate of –11.65 ±0.09 after application of 100 ton basalt ha^-1^ over 137 days. We presume that our unexpectedly high weathering rates are related to higher dissolved organic carbon (DOC) production due to the pasteurization process. During pasteurization, soil microbial necromass likely increased as a direct result of heating the soil at 80 °C for four hours. Pasteurization therefore resulted in an increase in DOC (Fig. S18), which has been shown to increase silicate weathering rates (Perez-Fodich and Derry, 2019).

In contrast with our hypothesis, AMF did not significantly increase basalt weathering rates. This outcome may be partly attributed to the experimental conditions: pasteurization likely increased nutrient availability (Hu et al., 2019) and, combined with fertilizer application (NPK; 96 – 10 – 39.5 kg ha^-1^), it ensured sufficient nutrient levels. Consequently, AMF might have played a limited role in basalt bio-weathering, as the experimental conditions may have reduced the need for AMF-driven nutrient mobilization from the mineral substrate.

#### 4.1.1 Reconciling contrasting weathering proxies

Pore water pH, alkalinity, Ca and Mg increased with basalt application, with no AMF effect, whereas pore water DIC showed a significant basalt x AMF x time interaction (Fig. 3-4). The observed increase in pore water DIC with basalt application was reinforced by AMF towards the end of the experiment, as the fungal hyphae developed. The observed increase in pore water DIC with AMF might suggest that AMF increased basalt weathering rates. However, other weathering proxies such as pore water pH, alkalinity, Ca and Mg did not show similar patterns. This discrepancy between DIC and the other weathering indicators may be due to different reasons.

While silicate weathering is expected to increase DIC through (bi)carbonate production, the observed increase in DIC may also reflect elevated soil partial CO₂ pressure (pCO₂). Pore water DIC is governed by carbonate equilibria, which are sensitive to transient environmental conditions such as pore water pH and pCO₂ (Hartmann et al., 2013; Zeebe and Wolf-Gladrow, 2001). Indeed, higher pCO_2_ can increase dissolved H_2_CO_3_ and therefore DIC. As the pore water pH in our experiment was > 6.5 in the second half of the experiment (Fig. 3), the dissolved H_2_CO_3_ likely speciated into (bi)carbonate ions within the soil solution, leading to elevated DIC.

A second possible explanation for the discrepancy between pore water DIC and alkalinity lies in the role of the anions in alkalinity. Total alkalinity is the sum of base cation charges, minus the sum of conservative anion charges (e.g. Cl*-*, SO ^2-^, PO_4_ ^3-^, NO *^-^*) (Barker, 2013; Wolf-Gladrow et al., 2007). While DIC increased with AMF, other anions may have decreased, leading to a null effect of AMF on alkalinity. We did not find a decrease in pore water NO *^-^*, SO ^2-^– and PO ^3-^ with AMF (Fig. 4; Fig. S3), but organic anions may still have contributed to the observed effect. During the middle of the growing season, basalt treatments without AMF tended to have lower DOC compared to basalt treatments with AMF (Fig. S18), partially supporting this hypothesis.

Finally, cation uptake may have also played a role. Plants and AMF act as a sink for cations, and a positive effect of AMF on this plant and AMF sink may have contributed to the observed decoupling between DIC and base cations. In this case, we would observe an increase in Ca and Mg stocks in the plant and/or AMF tissues. However, basalt only increased Mg stocks in plants, without an effect on Ca stocks (Fig. S19). Therefore, the plant sink hypothesis can only partially explain the observed mismatch between DIC and other weathering proxies such as pore water cations.

Although we cannot fully disentangle the divergence of different weathering proxies, the inconsistencies in their responses highlight the limitations of relying on a single proxy to assess weathering dynamics, as it may not adequately capture the complexity of the system (see also Iff et al. (2024)). According to the explicit conservative expression for total alkalinity (Wolf-Gladrow et al., 2007), DIC and alkalinity consist of distinct chemical species, thus complicating the interpretation of these weathering proxies. Therefore, our results suggest that cation accounting, rather than carbon accounting, may provide a more reliable estimate of weathering dynamics, as proposed by Bijma et al. (2025).

### 4.2 Basalt, but not AMF, improved soil fertility measures

Basalt, but not AMF, increased soil pH, CEC and base saturation (Fig. 2). The observed increases in soil pH, CEC, and base saturation upon basalt application align with our hypothesis of basalt improving soil fertility. The observed increase in base saturation was attributable to an increase in soil exchangeable Ca, Mg and Na due to basalt application (Fig. S20). Indeed, during weathering of basalt, protons are consumed and cations are released. Additionally, weathering of primary minerals promotes the formation of metal (hydr)oxides and secondary silicate phases, ultimately increasing the content of clay-sized particles in the soil and therefore CEC (Righi and Meunier, 1995). Our results corroborate previous work that observed increased soil pH and CEC upon basalt application to tropical soils (Anda et al., 2015; Conceição et al., 2022; Gillman, 1980). An increase in pH was also found in experiments conducted in using relatively younger, less cation-depleted soils upon basalt or dunite application (Rijnders et al., 2023; Rijnders et al., 2025; Skov et al., 2024; Ten Berge et al., 2012). These consistent results show that EW can contribute to soil restoration by counteracting soil acidification and improving CEC and availability of exchangeable bases, which are a common problem associated with intensive agriculture (Ashitha et al., 2021; Goulding, 2016).

#### 4.2.1 Limited AMF and basalt effects on soil nutrients

While basalt amendment increased soil CEC and pH, it generally did not affect soil nutrient concentrations, with a few exceptions. Despite an increase in exchangeable Ca and Mg, only Mg increased in the pore water as well. This might be at least partially explained by weathering kinetics: forsterite, a Mg-rich olivine, weathers relatively quickly compared to other Ca-bearing minerals within the applied basalt, such as labradorite, a Ca-enriched plagioclase feldspar (Gudbrandsson et al., 2011). Nevertheless, basalt increased exchangeable Ca (Fig. 6; Fig. S20), indicating that Ca was released during weathering, as observed by Li and Dong (2013), Niron et al. (2024) and Vienne et al. (2022). Our pore water and exchangeable Ca results also indicate that, once released, Ca was rapidly adsorbed onto soil exchange sites rather than remaining in solution.

The observed Mg and Ca increases in the exchangeable fraction are in agreement with existing literature, as application of basalt supplies new cations that can bind to soil particles, increasing base saturation (Gillman, 1980; Gillman et al., 2001, 2002; Te Pas et al., 2023). Increases in exchangeable Ca and Mg were also reported by Buss et al. (2024), whereas in a mesocosm experiment by Niron et al. (2024), basalt application did not affect any of the measured cations (Ca, Mg, K, Na, Fe and Al) in the exchangeable fraction, thus contrasting our findings. On the other hand, in our experiment exchangeable Zn, an important micronutrient, decreased with basalt application in the two top layers, a result consistent with previous research (Buss et al., 2024; Desmalles et al., 2025). Generally, above a pH of 6 exchangeable Zn contents tend to be very low (Blume et al., 2010). Therefore, the observed increase in soil pH from 5.65 ±0.11 to 6.36 ±0.17 following basalt application may have contributed to the reduction in exchangeable Zn.

Our finding of increased exchangeable Mg, Ca and Na (albeit not in all soil layers) is cautiously encouraging, given the global imperative to restore and maintain soil nutrient levels to support sustainable agricultural productivity (Un General Assembly, 2015). Indeed, the exchangeable pool consists of cations adsorbed onto soil particles that can be readily exchanged with the soil solution (Tessier et al., 1979), representing a key reservoir of nutrients, from which plant roots primarily acquire elements essential for growth (Roy et al., 2006). At the same time, even though basalt contains Fe, and in smaller amounts also K, which are important plant nutrients, their concentration in the exchangeable and pore water pools did not increase with basalt application, stressing the need to be cautious when expecting soil nutrient replenishment co-benefits from basalt application. Moreover, the observed decrease in exchangeable Zn underscores the need for careful evaluation of both unintended side-effects as well as potential co-benefits associated with enhanced weathering.

Finally, AMF did not significantly affect (micro)nutrient concentrations in the soil pore water nor in the soil exchangeable complex. Our results are thus in contrast with our hypotheses, as we expected increases in (micro)nutrient availability and/or uptake with AMF. This outcome may be at least partially attributed to the experimental conditions, namely the pasteurization process and the fertilizer addition, as explored at the end of section 4.1. In addition, the soil used in this experiment was relatively young and not particularly cation-depleted, therefore likely resulting in adequate nutrient availability, possibly masking any basalt– and/or AMF-driven effects on soil nutrients (Hu et al., 2019).

#### 4.2.2 Potential for future soil organic C stabilization and clay-sized particles formation

Most of the agriculturally relevant (micro)nutrients present in the basalt (Ca, Mg, Fe and Na) increased in the reducible pool in the 20-40 cm layer, and Ca and Si also increased in the oxidizable pool in the same layer (Table S7). The reducible and oxidizable pools comprise cations bound to metal (hydr)oxides and organic matter respectively (Tessier et al., 1979). Cations in these fractions are expected to be retained in the soil for a relatively long time, with plants able to access these nutrients only upon weathering (Niron et al., 2024; Palandri and Kharaka, 2004). Nevertheless, metal (hydr)oxide formation can ultimately increase CEC and govern SOC stabilization processes (Basile-Doelsch et al., 2015; Beerling et al., 2020; Manning et al., 2024; Totsche et al., 2017), with downstream benefits for plant growth and soil fertility, particularly in agricultural soils which are often depleted in organic matter (Rusco et al., 2001). These processes can improve soil structure, water retention, nutrient availability, and biological activity, thereby supporting overall soil health (Bot and Benites, 2005). Although the total stock of cations in the oxidizable fraction was relatively small compared to other soil pools, the observed increases in Ca and Si following basalt application suggest potential for organic matter stabilization with enhanced weathering.

### 4.3 Limited basalt and AMF effects on crop yield and quality

In line with our soil nutrient data and contrary to our hypotheses, plant nutrient concentrations were generally not affected by basalt application or AMF presence, with the exception of Mg, which generally increased with basalt. Increased plant Mg can be directly linked with the observed increase in pore water Mg concentrations and Mg exchange capacity, leading to elevated Mg uptake. Our results are supported by research carried out on corn after basalt application (Boniao et al., 2006; Rijnders et al., 2025), and on ryegrass (Ten Berge et al., 2012) and soybean (Moretti et al., 2019) after dunite application. These consistent findings indicate that EW can help alleviate Mg deficiency, which often occurs in agricultural systems that are fertilized with N, P, and K as uptake competition between these elements and Mg arise (Guo et al., 2016).

Nevertheless, our plant data shows only limited evidence for a positive effect of basalt and AMF on crop yield and quality, challenging the assumption that EW and AMF consistently deliver agronomic co-benefits. This outcome may be at least partially attributed to the experimental conditions: mycorrhizal symbiosis is established by the allocation of photosynthetic C from the plant to the fungus, and an important driver of this symbiosis is low bioavailability of soil P (Leake and Read, 2017). The sufficient P availability in our experiment (due to pasteurization and fertilization) might have led to lower energy investment in AMF production, hence possibly explaining the lack of basalt effect on AMF colonization and hyphal length.

In addition, the short experimental duration of our trial (113 days) might have limited the AMF effect on crop yield and quality. Indeed, a meta-analysis by Qin et al. (2022) identified experimental duration as the most important factor limiting the influence of AMF on crop growth, followed by soil texture, with a smaller AMF effect on sandy soils (>50% sand). Given the short experimental duration of the present experiment and texture of the used soil (sandy loam; 69.5% sand, 28.1% silt, 1.8% clay), it is likely that both factors played a role in the observed lack of AMF effect on plant biomass in our study.

The observed lack of a plant biomass response is similar to results by Vienne et al. (2022), where the application of 50 ton ha^-1^ of basalt did not significantly affect potato growth. In other experiments, on the other hand, basalt application increased biomass of spring oat (Skov et al., 2024), sorghum (Kelland et al., 2020), and maize (Rijnders et al., 2025). Additional counter evidence is provided by the review of Swoboda et al. (2022), which focused primarily on silicate rock powder applications in highly weathered, nutrient-poor soils, and which reported a positive silicate effect on crop biomass for almost all studies using mafic and ultramafic rocks as feedstock. These contrasting effects across soils and climatic regions emphasize the context-dependency of the effect of EW on crops.

### 4.4 Need for monitoring of toxic trace elements

Our findings highlight the need for careful monitoring of soil and plant heavy metal concentrations following basalt amendment. Given the observed increases in pore water Zn and Ni, our results do not support the hypothesis that basalt application does not elevate the availability of heavy metals in the soil. This is noteworthy, as Ni release is among the primary environmental safety concerns associated with EW in agricultural contexts (Suhrhoff, 2022; Te Pas et al., 2023). During the 113-day experimental period, average pore water Ni concentrations exceeded the freshwater Environmental Quality Standards (EQS) for both the EU (4 µg l⁻¹) and Australia (8 µg l⁻¹) in both control and basalt treatments (11.67 ±1.2 and 26.8 ±4.9 µg l⁻¹ respectively). This result might be partially due to background Ni present in the soil used for this experiment. Nevertheless, the observed concentrations remained below regulatory limits established by other jurisdictions, such as the United States (52 µg l⁻¹) and Canada (200 µg l⁻¹).

Plant heavy metal concentrations were generally unaffected by basalt amendment, with the notable exception of an increase in Ni in corn tissue (0.65 ±0.0004 and 0.42 ±0.0001 mg kg^-1^ for basalt +AMF and control –AMF respectively), therefore only partially supporting the hypothesis that basalt amendment does not increase plant heavy metal concentrations. Despite the increase, corn Ni concentrations remained below the EU regulatory threshold of 0.8 mg kg^-1^ which will come into effect in 2026 (European Commission, 2024). Our results therefore partially contrast with previous research, where Ni concentrations did not increase upon basalt application (Kelland et al., 2020; Rijnders et al., 2025; Vienne et al., 2022).

Heavy metal concentrations in the pore water and soil exchangeable complex were mostly not affected by AMF. On the other hand, AMF increased Cr concentration in the corn tissue. Therefore, our hypothesis that AMF presence would decrease heavy metal availability is not supported by our data. This finding underscores that AMF inoculation does not universally confer benefits and may, under certain conditions such as in co-deployment with silicate amendment, pose risks to food safety. It is important to note that Cr concentrations in corn tissue were, regardless of treatment, higher than the average Cr concentrations found in common vegetables as identified by WHO (World Health Organization, 1988), which is in the range of 0.005–0.03 mg Cr kg^-1^ fresh weight (Fig. S21).

## 5 Conclusion

Despite a synergistic effect of basalt and arbuscular mycorrhizal fungi (AMF) on pore water dissolved inorganic carbon (DIC), our 113-day mesocosm experiment did not reveal a significant AMF-induced enhancement of weathering rates, as assessed through cation mass balance, pore water alkalinity and pore water pH. These findings suggest that pore water DIC alone is not a reliable proxy for weathering dynamics in soil systems, given its sensitivity to a range of dynamic soil parameters such as partial CO_2_ pressure and pH. Notably, AMF had limited influence on nutrient availability, crop yield and quality under the fertilized conditions of this study. Longer-term experiments or those conducted under nutrient-limited conditions are needed to verify whether this lack of a short-term effect persists over time.

In contrast to AMF, basalt amendment improved several indicators of soil fertility, including soil pH, cation exchange capacity (CEC) and base saturation (particularly exchangeable Ca and Mg). These findings support the role of enhanced weathering in mitigating soil acidification and restoring base cation availability. However, these improvements did not translate into enhanced crop yield or nutrient status, with the exception of increased Mg concentrations in plant tissues. Therefore, our results indicate that the anticipated agronomic co-benefits of enhanced weathering are likely context-dependent and cannot be generalized to all soils and ecosystems. Moreover, although the increase of toxic trace elements in soil and plants was limited, our results stress the importance of carefully evaluating the environmental safety of enhanced weathering for the soil under consideration.

## Code and data availability

The data and model code in support of our findings are openly available in Zenodo at https://doi.org/10.5281/zenodo.16813184 (Boito et al., 2025).

## Author contributions

LB and JR contributed equally to this manuscript. LB and JR: conceptualization, data curation, formal analysis, investigation, project administration, software, supervision, visualization, writing – original draft, writing – review and editing. LS: conceptualization, data curation, formal analysis, funding acquisition, investigation, software, methodology, project administration, supervision, writing – review and editing. PF: investigation, methodology, writing – review and editing. MM: investigation, writing – review and editing. AV: conceptualization, formal analysis, funding acquisition, software, writing – review and editing. EV: methodology, writing – review and editing. SV: conceptualization, formal analysis, funding acquisition, methodology, project administration, resources, supervision, writing – review and editing.

## Acknowledgements

We thank Emma Pellegrini, Martín “Gato” Carrera Larrea, Sarah Janse and Jasper Roussard for their invaluable contributions in data collection. We thank the Helmholtz Laboratory for the Geochemistry of the Earth Surface (HELGES) for carrying out elemental analyses of the porewater and the sequential extraction protocol of soil samples.

