## Supplementary Information for "Effects of basalt amendment and mycorrhizal inoculation on soil chemical properties and maize growth"

Supplementary information relative to the manuscript titled “Effects of basalt amendment and mycorrhizal inoculation on soil chemical properties and maize growth”

Lucilla Boito, Jet Rijnders, Laura Steinwidder, Patrick Frings, Arthur Vienne, Mirthe Maes, Erik Verbruggen, and Sara Vicca

Figure S1: Particle size distribution of basalt. P80= 80% of the particles having a diameter less than or equal to this size.

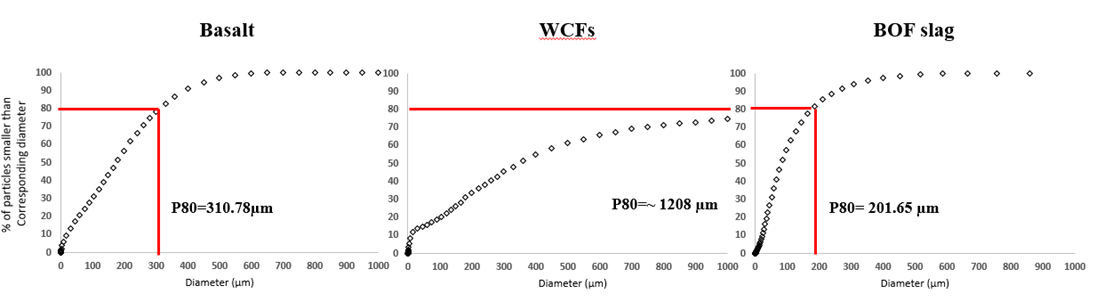

Table S2: Dry weights of corn smut galls for each treatment (C-AMF = control without AMF, C+AMF = control with AMF, B-AMF = basalt without AMF, B+AMF = basalt with AMF) at the end of the experiment. p- and F-values are shown from a general linear model with gall weight as response variable and basalt, AMF and their interaction as covariables. The interaction was not significant (ns) and was excluded from the model.

|  | Fungus dry weight (g) | | | | Statistical analysis | | |
| --- | --- | --- | --- | --- | --- | --- | --- |
| *Replica* | **B+AMF** | **B-AMF** | **C+AMF** | **C-AMF** | **Variable** | **p-value** | **F-value** |
| *1* | 0 | 0 | 0 | 0 | Basalt | 0.29 | 1.19 |
| *2* | 0.27 | 0 | 0 | 0.06 | AMF | 0.93 | 0.05 |
| *3* | 0.51 | 0.06 | 0.21 | 0.27 | Basalt^AMF | ns | ns |
| *4* | 0.53 | 1.38 | 0.35 | 0.44 |  |  |  |
| *5* | 6.89 | 6.96 | 2.05 | 0.67 |  |  |  |

Figure S3: Mean concentrations of copper (Cu), manganese (Mn), ammonium-nitrogen (NH_4_^+^-N), nitrate-nitrogen (NO_3_^-^-N) and sulphate (SO_4_^2-^) in the porewater during the experiment for the four treatments (C-AMF = control without AMF, C+AMF = control with AMF, B-AMF = basalt without AMF, B+AMF = basalt with AMF). Error bars represent the standard error on the mean. p- and F-values are shown from a linear regression analysis with elements as response variables and basalt, AMF and time, and their relevant interactions, as covariables. When interactions were not significant, they were excluded from the model and not shown here. Interactions are indicated by carets (^), while significant relationships by an asterisk (*).

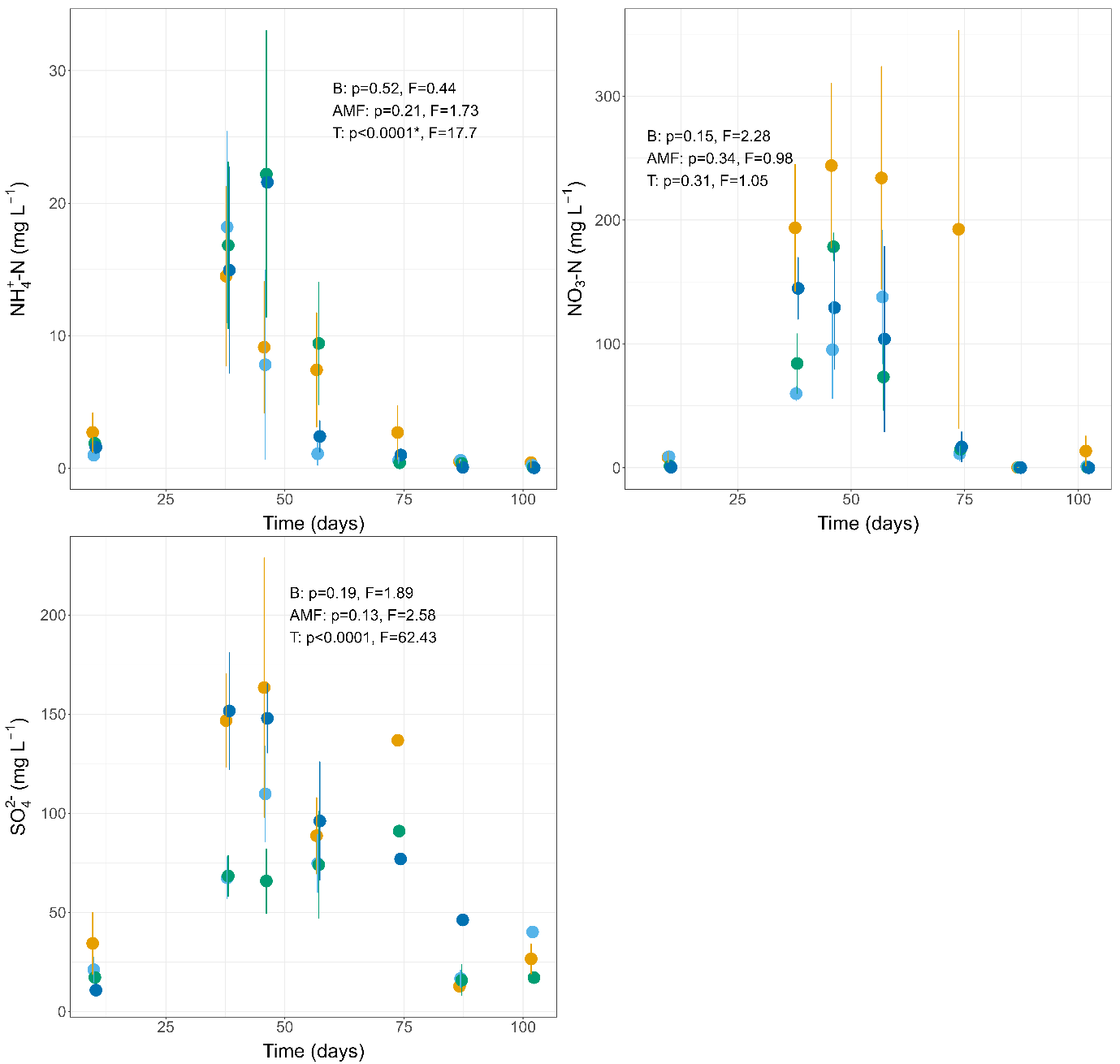

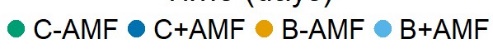

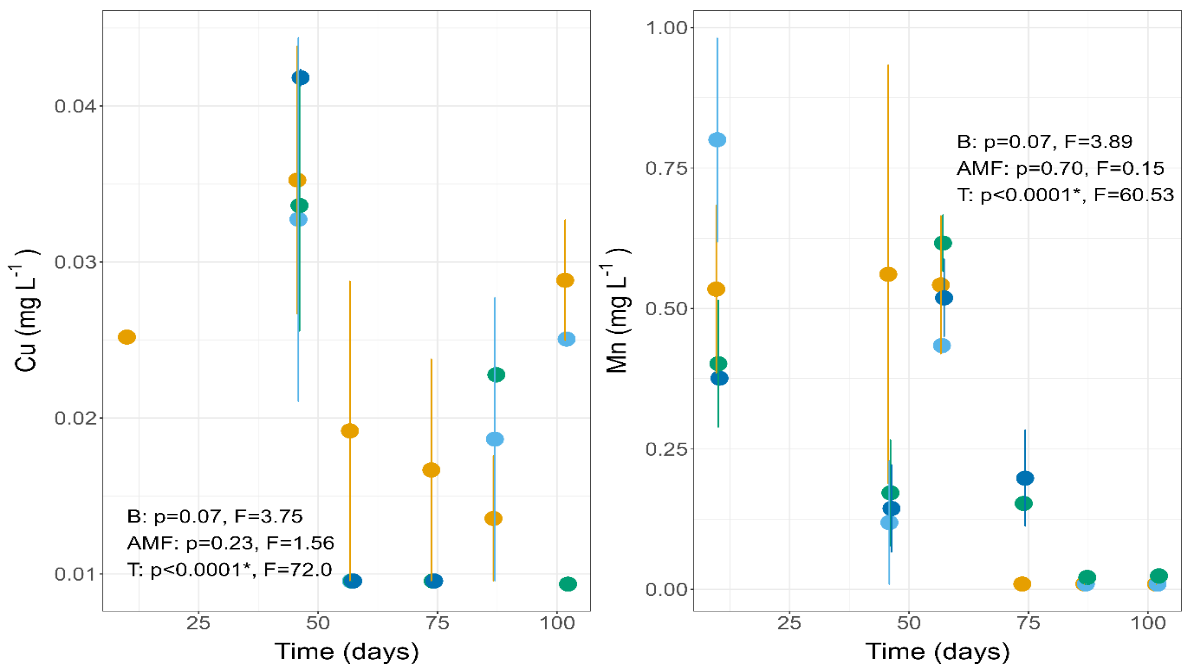

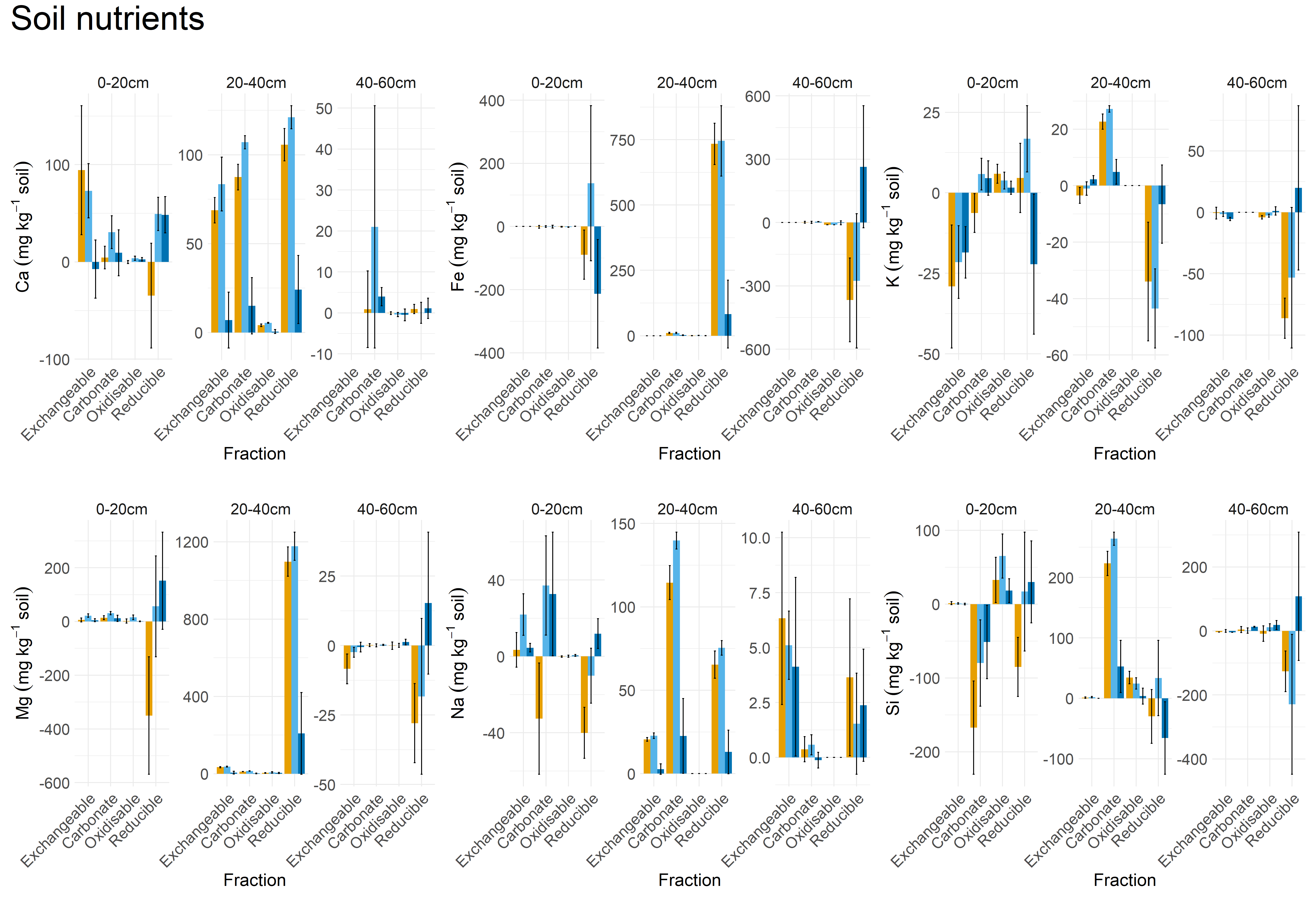

Figure S4: Mean soil nutrient concentrations (Ca, Fe, K, Mg, Na and Si) for the three treatments (B-AMF = basalt without AMF, B+AMF = basalt with AMF, C+AMF = control with AMF) at the end of the growing season. Nutrient concentrations are shown as the difference between the start and the end of the growing season, and are shown compared to the control treatment (control without AMF). Soils were divided into three sampling depths (0-20 cm, 20-40 cm, 40-60 cm) for which nutrient concentrations were measured in four soil fractions (exchangeable, carbonate, oxidisable and reducible fraction) with sequential extraction analysis according to Tessier et al (1979).

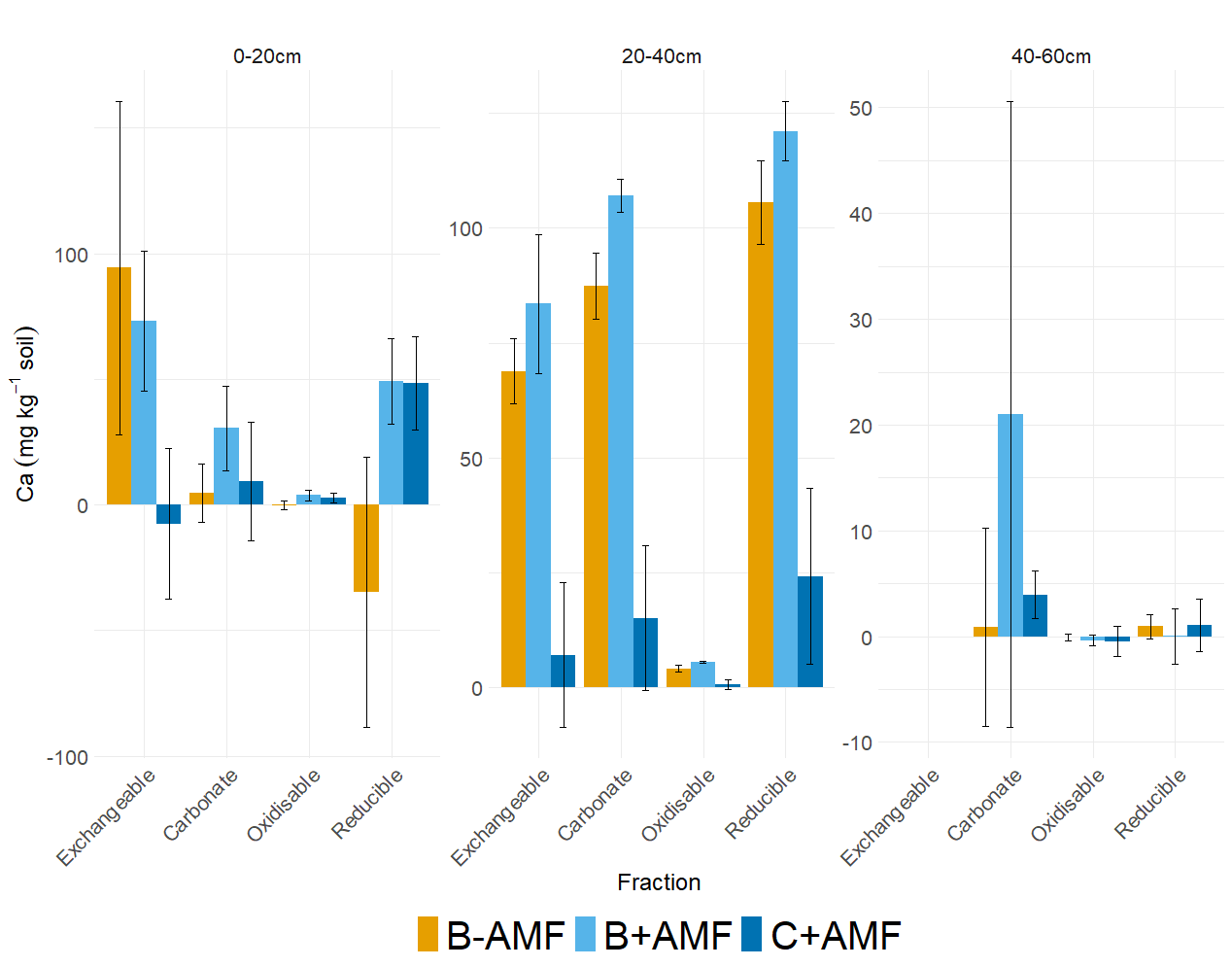

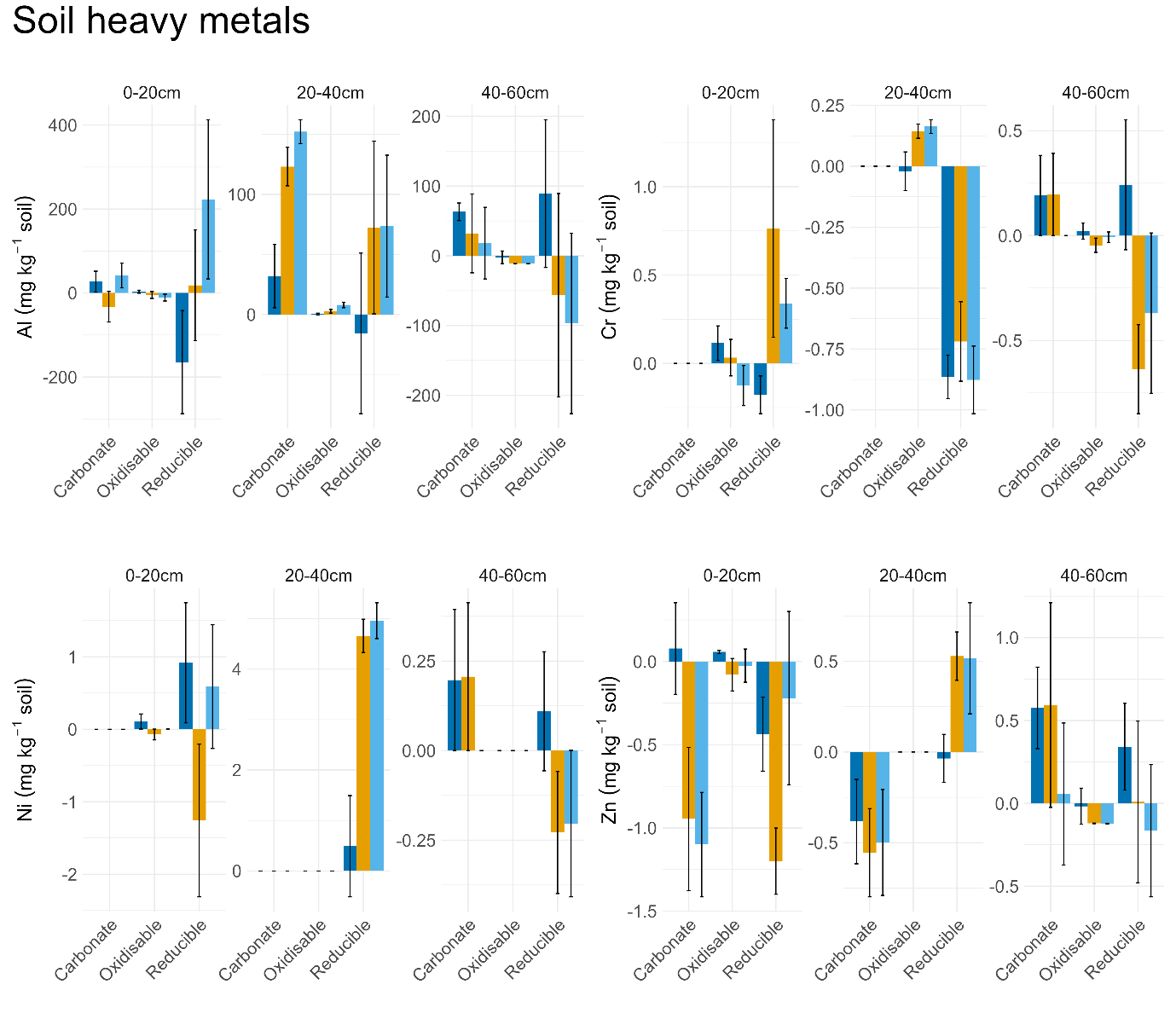

Figure S5: Mean soil heavy metals (Al, Cr, Ni and Zn) concentrations for the three treatments (B-AMF = basalt without AMF, B+AMF = basalt with AMF, C+AMF = control with AMF) at the end of the growing season. Concentrations are shown as the difference between the start and the end of the growing season, and are shown compared to the control treatment (control without AMF). Soils were divided into three sampling depths (0-20 cm, 20-40 cm, 40-60 cm) for which nutrient concentrations were measured in four soil fractions (exchangeable, carbonate, oxidisable and reducible fraction) with the sequential extraction method after Tessier et al. (1979).

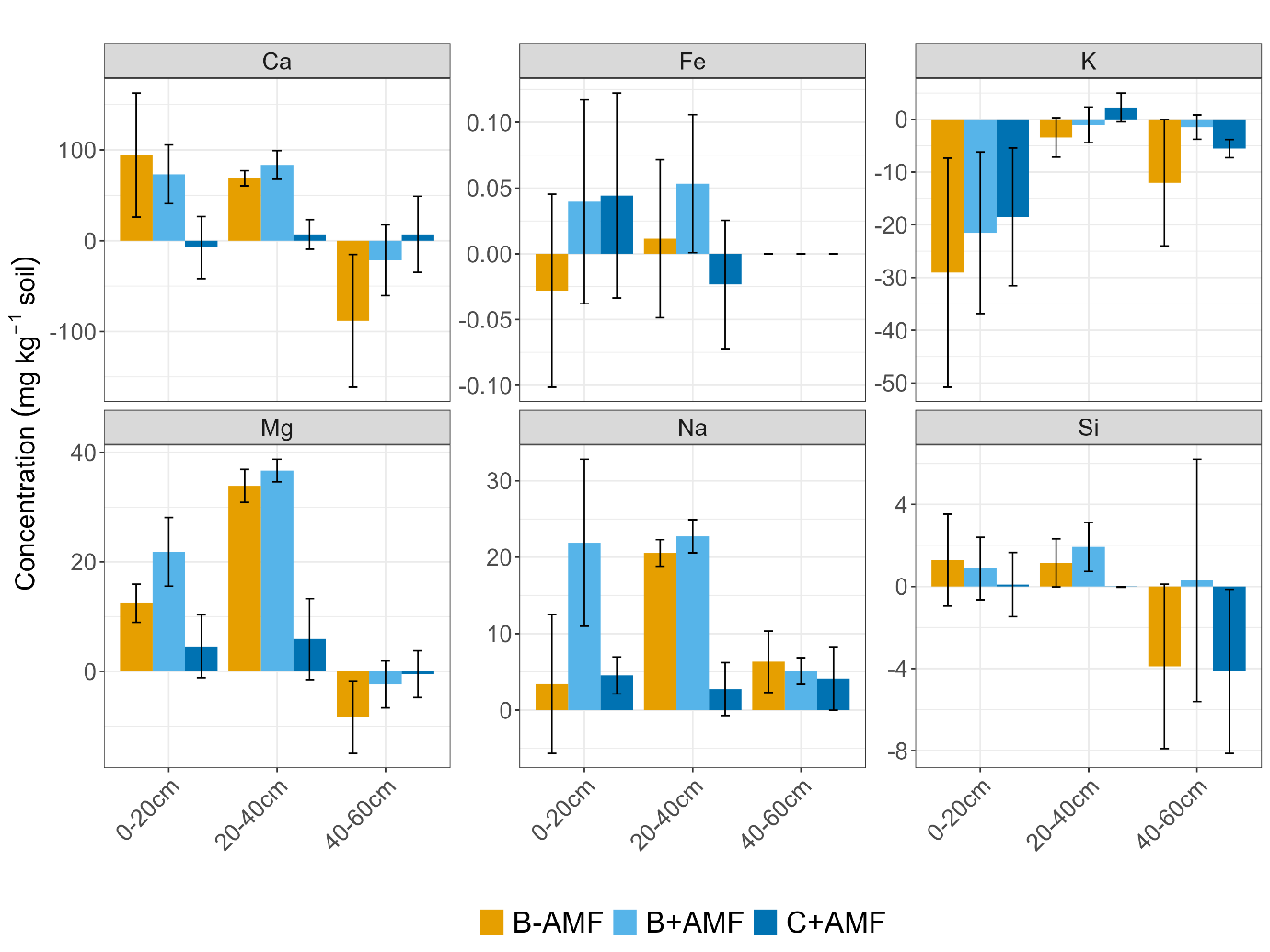

Table S6: p- and F-values from a linear regression analysis with element concentration in four soil fractions and for three different soil layers as response variable, and basalt, AMF and their interaction as covariates. Interactions that were not significant were excluded from the model. Significant relationships are indicated in bold, while p values between 0.05 and 0.099 in italics. Whenever assumptions of normality or homoscedasticity were not met, the non-parametric Kruskal-Wallis test was used instead, with treatment as covariate. In those cases, p-and chi² values are shown. Whenever element concentrations for any fraction-layer combination were below detection (bd) or below the limit of quantification (<LOQ), statistical analysis was not possible.

|  |  |  | Basalt | | AMF | | Basalt ^ AMF | |
| --- | --- | --- | --- | --- | --- | --- | --- | --- |
| Element | Layer | Fraction | p | F | p | F | p | F |
| Calcium (Ca) | 0-20 | Carbonate | 0.41 | 0.71 | 0.28 | 1.25 | ns | ns |
|  |  | Exchangeable | **0.037** | **5.12** | 0.72 | 0.14 | ns | ns |
|  |  | Oxidisable | 0.97 | 0.001 | *0.095* | *3.1* | ns | ns |
|  |  | Reducible | Kruskal-Wallis test | | p=0.49; chi^2^=2.45 | |  |  |
|  | 20-40 | Carbonate | **Kruskal-Wallis test** | | **p<0.001; chi²=16.63** | |  |  |
|  |  | Exchangeable | **<0.0001** | **40.77** | 0.35 | 0.9 | ns | ns |
|  |  | Oxidisable | **<0.0001** | **43.63** | 0.18 | 1.95 | ns | ns |
|  |  | Reducible | **Kruskal-Wallis test** | | **p<0.01; chi²=14.96** | |  |  |
|  | 40-60 | Carbonate | Kruskal-Wallis test | | p=0.72; chi²=1.35 | |  |  |
|  |  | Exchangeable | 0.23 | 1.58 | 0.76 | 0.09 | ns | ns |
|  |  | Oxidisable | Kruskal-Wallis test | | p=0.66; chi²=1.62 | |  |  |
|  |  | Reducible | 0.95 | <0.01 | 0.9 | 0.02 | ns | ns |
| Magnesium (Mg) | 0-20 | Carbonate | **0.017** | **6.98** | *0.053* | *4.3* | ns | ns |
|  |  | Exchangeable | **<0.01** | **11.58** | 0.15 | 2.29 | ns | ns |
|  |  | Oxidisable | 0.3 | 1.4 | 0.28 | 1.23 | ns | ns |
|  |  | Reducible | Kruskal-Wallis test | | p=0.21; chi^2^=4.53 | | ns | ns |
|  | 20-40 | Carbonate | **Kruskal-Wallis test** | | **p<0.01; chi²= 16.2** | |  |  |
|  |  | Exchangeable | **Kruskal-Wallis test** | | **p<0.01; chi²=12.3** | |  |  |
|  |  | Oxidisable | 0.15 | 2.27 | 0.17 | 2.06 | ns | ns |
|  |  | Reducible | **Kruskal-Wallis test** | | **p<0.01; chi²= 14.39** | | ns | ns |
|  | 40-60 | Carbonate | 0.35 | 0.92 | *0.07* | 3.71 | ns | ns |
|  |  | Exchangeable | 0.45 | 0.59 | 0.94 | <0.01 | ns | ns |
|  |  | Oxidisable | 0.54 | 0.4 | 0.57 | 0.34 | ns | ns |
|  |  | Reducible | 0.14 | 2.41 | 0.84 | 0.04 | ns | ns |
| Sodium (Na) | 0-20 | Carbonate | 0.6 | 0.28 | **0.048** | **4.5** | ns | ns |
|  |  | Exchangeable | **Kruskal-Wallis test** | | **p=0.046; chi^2^=7.98** | | ns | ns |
|  |  | Oxidisable | 0.52 | 0.43 | 0.42 | 0.7 | ns | ns |
|  |  | Reducible | **0.006** | 9.4 | *0.059* | *4.07* | ns | ns |
|  | 20-40 | Carbonate | **Kruskal-Wallis test** | | **p<0.01; chi²= 15.2** | |  |  |
|  |  | Exchangeable | **<0.0001** | 83.5 | 0.27 | 1.29 | ns | ns |
|  |  | Oxidisable | bd | | | | | |
|  |  | Reducible | **Kruskal-Wallis test** | | **p<0.01; chi²= 13.8** | | ns | ns |
|  | 40-60 | Carbonate | 0.23 | 1.53 | 0.95 | <0.01 | ns | ns |
|  |  | Exchangeable | Kruskal-Wallis test | | p=0.13; chi^2^=5.64 | | ns | ns |
|  |  | Oxidisable | bd | | | | | |
|  |  | Reducible | 0.55 | 0.37 | 0.92 | 0.01 | ns | ns |
| Potassium (K) | 0-20 | Carbonate | 0.63 | 0.24 | *0.08* | *3.49* | ns | ns |
|  |  | Exchangeable | 0.23 | 1.55 | 0.68 | 0.18 | ns | ns |
|  |  | Oxidisable | 0.22 | 1.61 | 0.55 | 0.37 | ns | ns |
|  |  | Reducible | 0.15 | 2.32 | 0.7 | 0.15 | ns | ns |

Table S6: continued.

|  |  |  | Basalt | | AMF | | Basalt ^ AMF | |
| --- | --- | --- | --- | --- | --- | --- | --- | --- |
| Element | Layer | Fraction | p | F | p | F | p | F |
| Potassium (K) | 20-40 | Carbonate | **Kruskal-Wallis test** | | **p<0.001; chi²=17** | |  |  |
|  |  | Exchangeable | 0.15 | 2.27 | 0.31 | 1.08 | ns | ns |
|  |  | Oxidisable | bd | | | | | |
|  |  | Reducible | **0.02** | **6.57** | 0.45 | 0.58 | ns | ns |
|  | 40-60 | Carbonate | bd | | | | | |
|  |  | Exchangeable | 0.45 | 0.6 | 0.23 | 1.57 | ns | ns |
|  |  | Oxidisable | 0.36 | 0.88 | 0.34 | 0.97 | ns | ns |
|  |  | Reducible | 0.09 | 3.21 | 0.86 | 0.03 | ns | ns |
| Iron (Fe) | 0-20 | Carbonate | 0.99 | <0.01 | 0.83 | 0.05 | ns | ns |
|  |  | Exchangeable | 0.76 | 0.1 | 0.26 | 1.35 | ns | ns |
|  |  | Oxidisable | **0.03** | **5.87** | 0.56 | 0.36 | ns | ns |
|  |  | Reducible | 0.54 | 0.39 | 0.95 | 0.003 | ns | ns |
|  | 20-40 | Carbonate | **<0.0001** | **32.1** | 0.54 | 0.39 | ns | ns |
|  |  | Exchangeable | 0.29 | 1.18 | 0.82 | 0.05 | ns | ns |
|  |  | Oxidisable | *0.08* | 3.4 | 0.27 | 1.31 | ns | ns |
|  |  | Reducible | **<0.0001** | **35.1** | 0.89 | 0.02 | ns | ns |
|  | 40-60 | Carbonate | Kruskal-Wallis test | | p=0.92; chi²=0.49 | |  |  |
|  |  | Exchangeable | bd | | | | | |
|  |  | Oxidisable | Kruskal-Wallis test | | p=0.501; chi²=2.36 | | ns | ns |
|  |  | Reducible | *0.09* | *3.18* | 0.91 | 0.01 | ns | ns |
| Silicon (Si) | 0-20 | Carbonate | Kruskal-Wallis test | | p=0.29; chi²=3.76 | |  |  |
|  |  | Exchangeable | Kruskal-Wallis test | | p=0.6; chi²=1.88 | |  |  |
|  |  | Oxidisable | 0.2 | 1.79 | 0.44 | 0.62 | ns | ns |
|  |  | Reducible | 0.43 | 0.66 | 0.28 | 1.22 | ns | ns |
|  | 20-40 | Carbonate | **Kruskal-Wallis test** | | **p<0.01; chi²=15.1** | |  |  |
|  |  | Exchangeable | Kruskal-Wallis test | | p=0.5; chi²=2.39 | |  |  |
|  |  | Oxidisable | **<0.01** | **8.53** | 0.56 | 0.36 | ns | ns |
|  |  | Reducible | 0.22 | 1.62 | 0.27 | 1.26 | ns | ns |
|  | 40-60 | Carbonate | Kruskal-Wallis test | | p=0.52; chi²=2.28 | |  |  |
|  |  | Exchangeable | 0.15 | 2.31 | 0.13 | 2.57 |  |  |
|  |  | Oxidisable | 0.7 | 0.16 | 0.41 | 0.72 | ns | ns |
|  |  | Reducible | 0.13 | 2.5 | 0.89 | 0.02 | ns | ns |
| Aluminum (Al) | 0-20 | Carbonate | 0.77 | 0.09 | *0.07* | *3.81* | ns | ns |
|  |  | Exchangeable | 0.55 | 0.37 | 0.13 | 2.53 | ns | ns |
|  |  | Oxidisable | 0.15 | 2.28 | 0.77 | 0.09 | ns | ns |
|  |  | Reducible | 0.17 | 2.04 | 0.92 | 0.01 | ns | ns |
|  | 20-40 | Carbonate | **<0.0001** | **31.7** | 0.14 | 2.42 | ns | ns |
|  |  | Exchangeable | 0.22 | 1.63 | 0.87 | 0.03 | ns | ns |
|  |  | Oxidisable | **<0.01** | **10.13** | *0.099* | *3.04* | ns | ns |
|  |  | Reducible | 0.24 | 1.47 | 0.75 | 0.11 | ns | ns |
|  | 40-60 | Carbonate | Kruskal-Wallis test | | p=0.79; chi²=1.04 | |  |  |
|  |  | Exchangeable | bd | | | | | |
|  |  | Oxidisable | Kruskal-Wallis test | | p=0.5; chi²=2.36 | |  |  |
|  |  | Reducible | 0.25 | 1.43 | 0.97 | <0.01 | ns |  |

Table S6: continued.

|  |  |  | Basalt | | AMF | | Basalt ^ AMF | |
| --- | --- | --- | --- | --- | --- | --- | --- | --- |
| Element | Layer | Fraction | p | F | p | F | p | F |
| Cromium (Cr) | 0-20 | Carbonate | <LOQ | | | | | |
|  |  | Exchangeable | <LOQ | | | | | |
|  |  | Oxidisable | Kruskal-Wallis test | | p=0.15; chi²=5.35 | |  |  |
|  |  | Reducible | **0.049** | **4.47** | 0.41 | 0.71 | ns | ns |
|  | 20-40 | Carbonate | <LOQ | | | | | |
|  |  | Exchangeable |  |  |  |  |  |  |
|  |  | Oxidisable | *Kruskal-Wallis test* | | *p=0.09; chi^2^= 6.43* | |  |  |
|  |  | Reducible | 0.13 | 2.51 | *0.08* | *3.57* | ns | ns |
|  | 40-60 | Carbonate | Kruskal-Wallis test | | p=0.55; chi^2^= 2.12 | |  |  |
|  |  | Exchangeable | bd | | | | | |
|  |  | Oxidisable | 0.36 | 0.87 | 0.48 | 0.54 | ns | ns |
|  |  | Reducible | *0.055* | *4.23* | 0.8 | 0.07 | ns | ns |
| Nickel (Ni) | 0-20 | Carbonate | <LOQ | | | | | |
|  |  | Exchangeable | <LOQ | | | | | |
|  |  | Oxidisable | <LOQ | | | | | |
|  |  | Reducible | 0.32 | 1.05 | *0.09* | *3.2* | ns | ns |
|  | 20-40 | Carbonate | <LOQ | | | | | |
|  |  | Exchangeable | <LOQ | | | | | |
|  |  | Oxidisable | bd | | | | | |
|  |  | Reducible | **Kruskal-Wallis test** | | **p<0.01; chi² = 14.4** | |  |  |
|  | 40-60 | Carbonate | Kruskal-Wallis test | | p=0.55; chi^2^ = 2.12 | |  |  |
|  |  | Exchangeable | bd | | | | | |
|  |  | Oxidisable | bd | | | | | |
|  |  | Reducible | 0.11 | 2.91 | 0.92 | 0.01 | ns | ns |
| Zinc (Zn) | 0-20 | Carbonate | **<0.01** | **13.93** | 0.9 | 0.01 | ns | ns |
|  |  | Exchangeable | **<0.001** | **20.3** | 0.51 | 0.46 | ns | ns |
|  |  | Oxidisable | Kruskal-Wallis test | | p=0.55; chi^2^ = 2.09 | |  |  |
|  |  | Reducible | *0.07* | *3.76* | 0.46 | 0.57 | *0.07* | 3.74 |
|  | 20-40 | Carbonate | *Kruskal-Wallis test* | | *p=0.09; chi²= 6.38* | |  |  |
|  |  | Exchangeable | **Kruskal-Wallis test** | | **p<0.01*; chi² =11.66** | |  |  |
|  |  | Oxidisable | bd | | | | | |
|  |  | Reducible | **0.045** | **4.65** | 0.65 | 0.22 | ns | ns |
|  | 40-60 | Carbonate | 0.66 | 0.2 | 0.76 | 0.09 | ns | ns |
|  |  | Exchangeable | Kruskal-Wallis test | | p=0.28; chi^2^= 3.8 | |  |  |
|  |  | Oxidisable | Kruskal-Wallis test | | p=0.44; chi^2^= 2.69 | |  |  |
|  |  | Reducible | 0.38 | 0.8 | 0.98 | <0.01 | ns | ns |

Table S7: Direction of the basalt effect on elements in the reducible and oxidizable pools of two soil layers (elements in the 40-60 cm layer were not affected by basalt amendment in the reducible and oxidizable pool) at the end of the experiment. A “+” symbolizes a basalt-driven increase in elemental concentration, while a “-“ symbolizes a decrease. Only significant effects are reported. Trends are indicated in italics. No significant AMF effect was detected.

| Element | 0-20 cm layer | | 20-40 cm layer | |
| --- | --- | --- | --- | --- |
|  | Reducible | Oxidizable | Reducible | Oxidizable |
| Ca |  |  | + | + |
| Mg |  |  | + |  |
| Fe |  |  | + | - |
| Na |  |  | + |  |
| K |  |  | - |  |
| Si |  |  |  | + |
| Cr |  |  |  | *+* |
| Ni |  |  |  |  |
| Zn |  |  |  |  |
| Al |  |  |  | + |

Table S8: Leachate volumes during the experiment for each mesocosm. Only on two dates at the beginning of the growing season, leachates were collected. There were no leachates after 20 June 2022.

| Mesocosm | Treatment | Date | Volume (L) |
| --- | --- | --- | --- |
| 1 | B+AMF | 2022-06-10 | 1.7 |
|  |  | 2022-06-20 | 0.1 |
| 5 | B-AMF | 2022-06-10 | 0.88 |
| 10 | B+AMF | 2022-06-10 | 0.39 |
|  |  | 2022-06-20 | 0.16 |
| 14 | B-AMF | 2022-06-10 | 0.81 |
|  |  | 2022-06-20 | 0.09 |
| 18 | C+AMF | 2022-06-10 | 0.1 |
| 22 | C-AMF | 2022-06-20 | 0.1 |
| 25 | C+AMF | 2022-06-10 | 0.52 |
| 28 | C+AMF | 2022-06-10 | 0.076 |
| 37 | B-AMF | 2022-06-10 | 0.85 |
|  |  | 2022-06-20 | 0.05 |
| 42 | C+AMF | 2022-06-10 | 0.53 |
| 46 | C-AMF | 2022-06-10 | 0.79 |
| 50 | B+AMF | 2022-06-10 | 0.87 |
|  |  | 2022-06-20 | 0.095 |
| 53 | B+AMF | 2022-06-10 | 0.63 |
|  |  | 2022-06-20 | 0.3 |
| 56 | C-AMF | 2022-06-10 | 0.06 |
|  |  | 2022-06-10 | 0.222 |
| 62 | B-AMF | 2022-06-10 | 0.7 |

Figure S9: Concentrations of C, N, C/N ratio, Ca, Fe, K, P, Si, Al, Cd, Pb, V and Zn in the corn, leaves, stem, tassel, and roots for the four treatments (C-AMF = control without AMF, C+AMF = control with AMF, B-AMF = basalt without AMF, B+AMF = basalt with AMF). C and N content were determined by dry combustion based on the Dumas method using an elemental analyser (FLASH 2000, Interscience, Louvain-la-Neuve, Belgium). For each plant sample, 0.3 g was weighed and digested with HNO3 and H2O2 according to Walinga et al. (1989). Corn and leaf V concentrations were all below LOQ (<0.0001 mg g-1) and therefore are not presented. Boxes represent the interquartile range (25th–75th percentile), horizontal lines indicate the median, and whiskers extend to the most extreme data points within 1.5× of the interquartile range. Points outside this range are plotted as outliers. Note that for Al, Cd, Fe, Pb and V, the primary axis shows the concentration in the corn, leaves, stems, and tassels while the secondary axis shows the concentration in the roots. p- and F-values are shown in table S17. Significant differences are indicated with a different letter, while a similar letter means no differences between those treatments. Black letters indicate basalt effects, while red letter indicate AMF effects. If no letters are shown for a combination of plant part and element, this means that there was no significant influence of basalt or AMF.

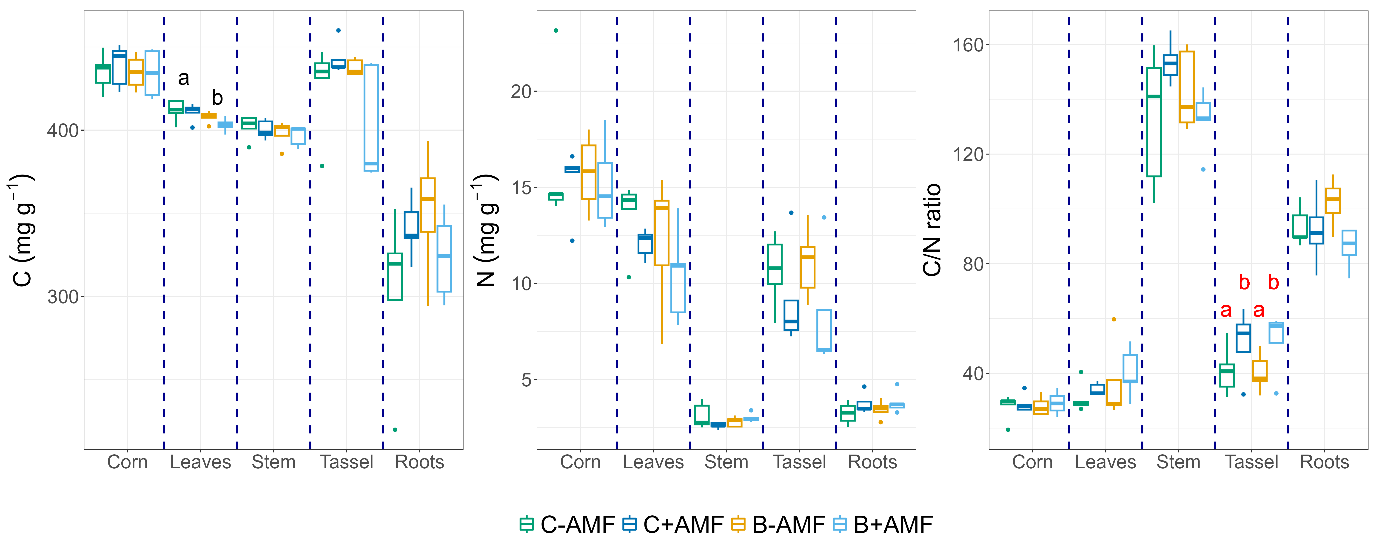

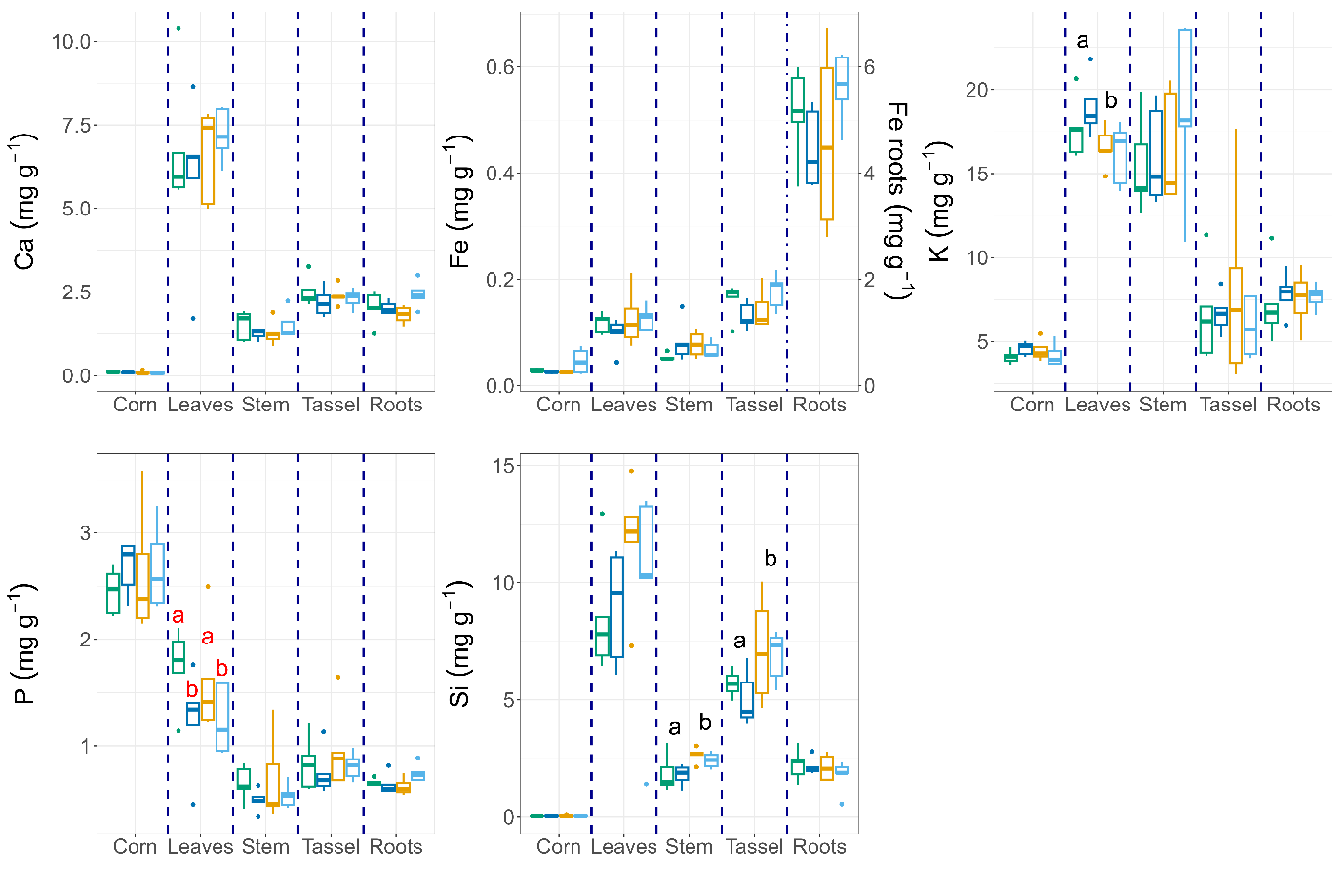

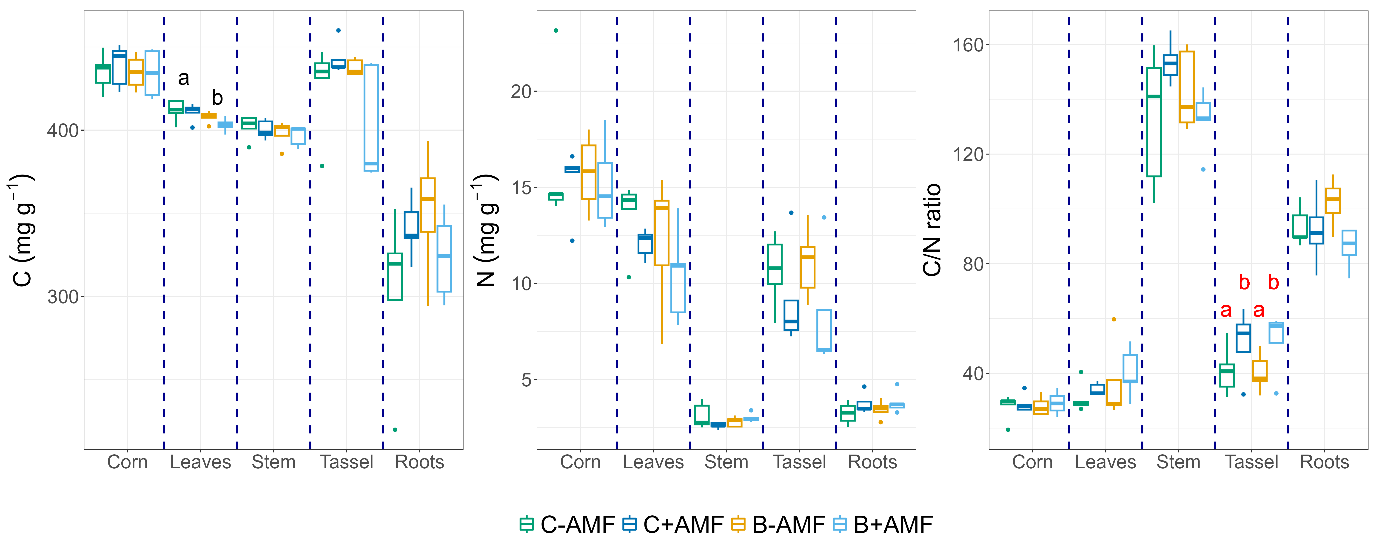

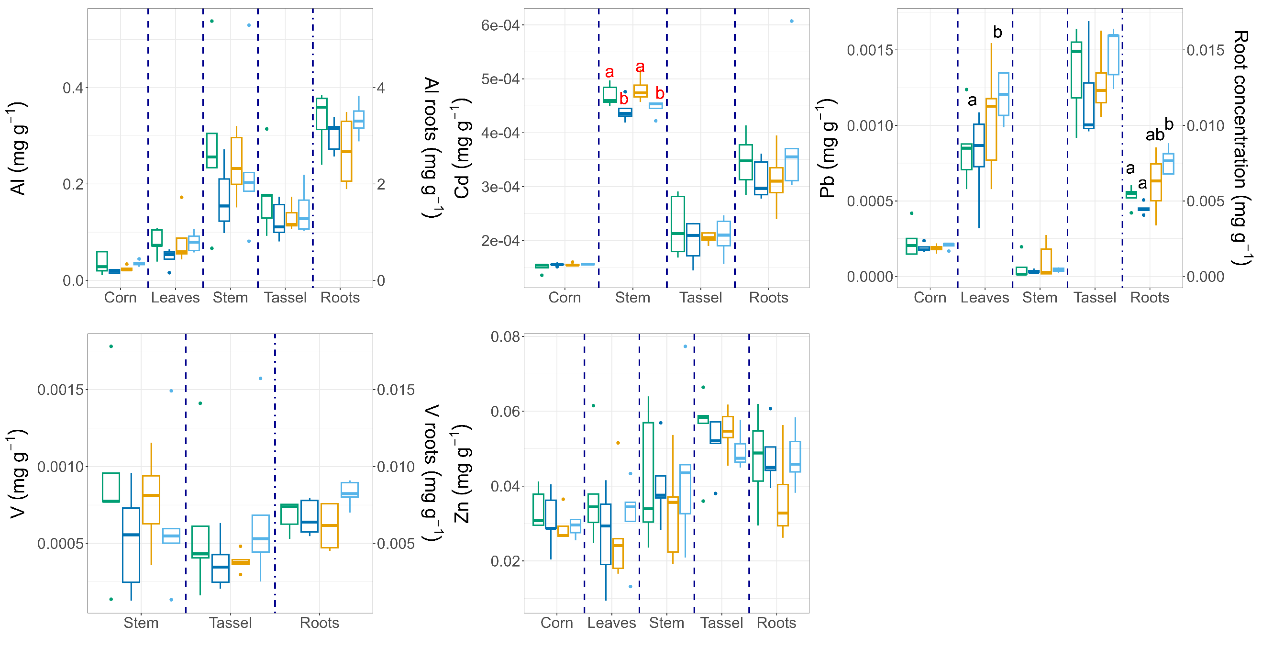

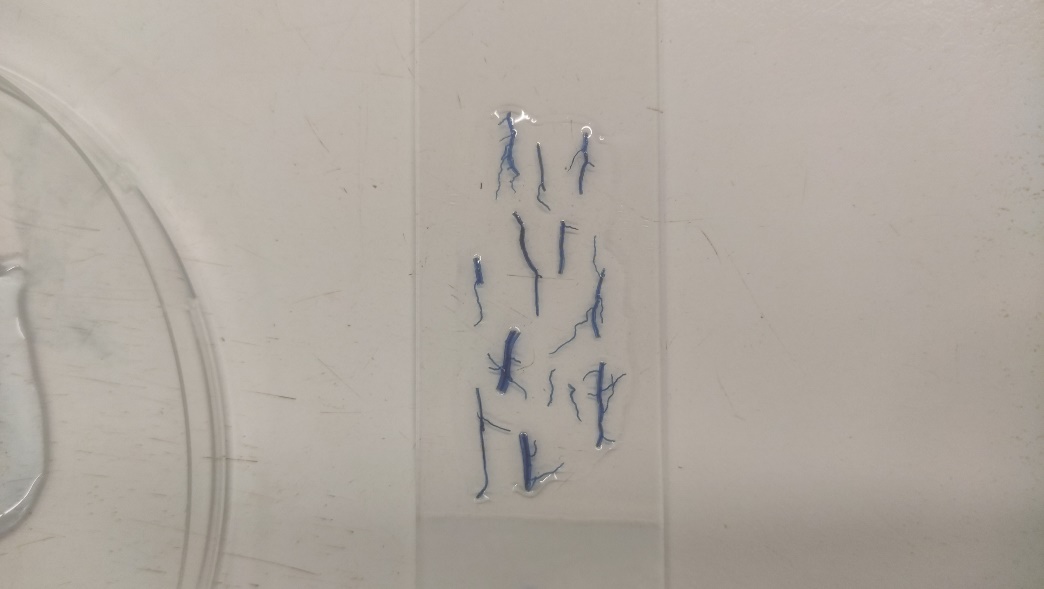

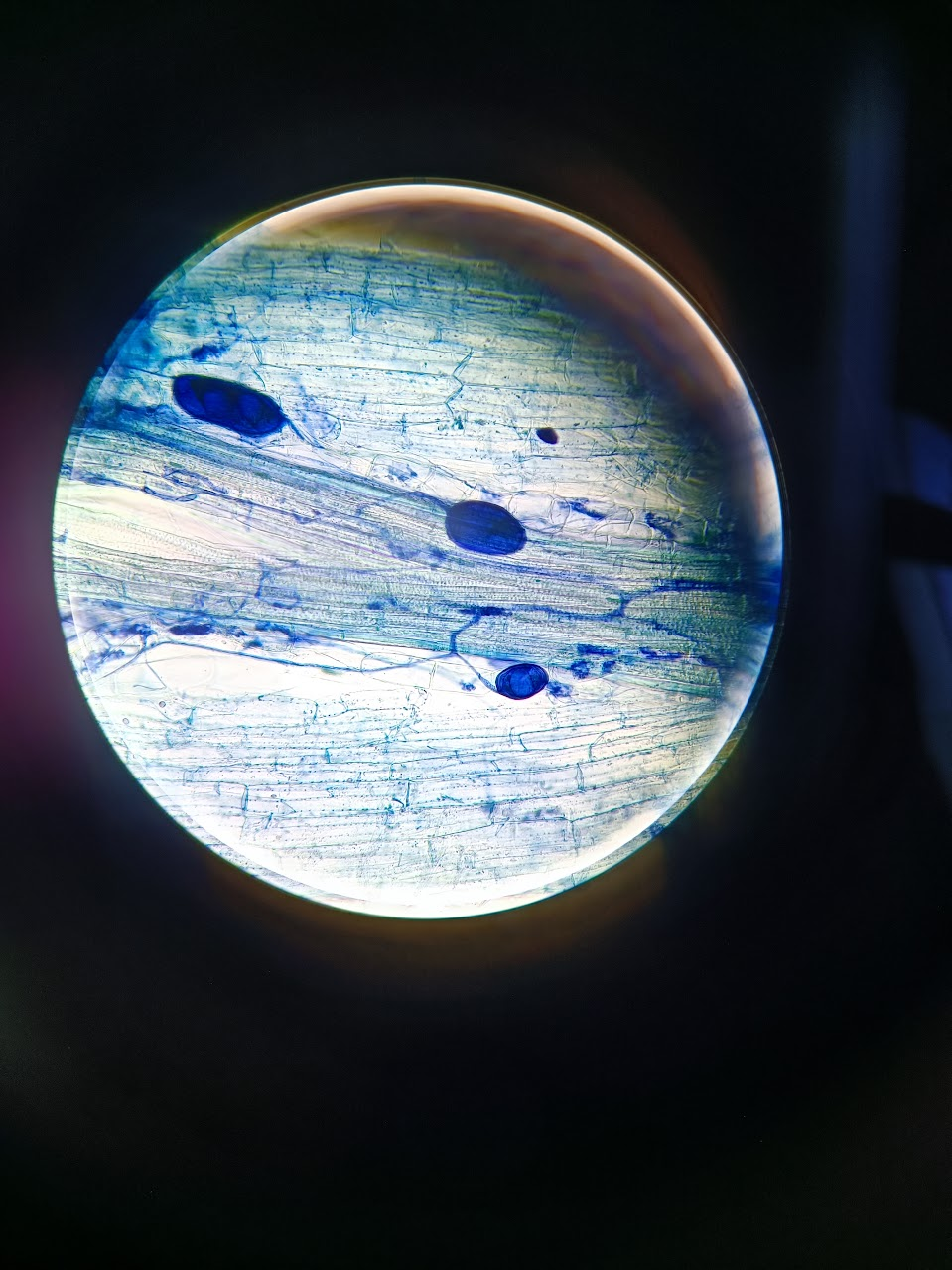

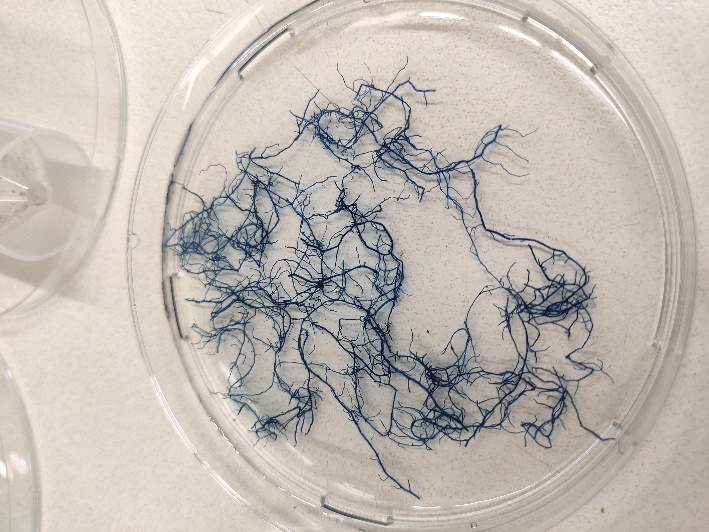

Figure S10: Overview of AMF colonization analysis. a) Roots stained with ink (10% Shaeffer Black ink in 10% acetic acid) b) Slide with stained roots, c) Root under microscope with vesicles, arbuscules and hyphae.

Arbuscule

Vesicle

b

a

C

Hyphae

Figure S11: Overview of AMF hyphal length analysis. a) harvested mesh bag, b) Set-up of the vacuum system where a vacuum pump is connected to the outlet to remove the liquid from the filter, which is placed in between the reservoir and the tube on top, c) a slide is prepared with the filter to quantify the AMF hyphal length under an optical microscope.

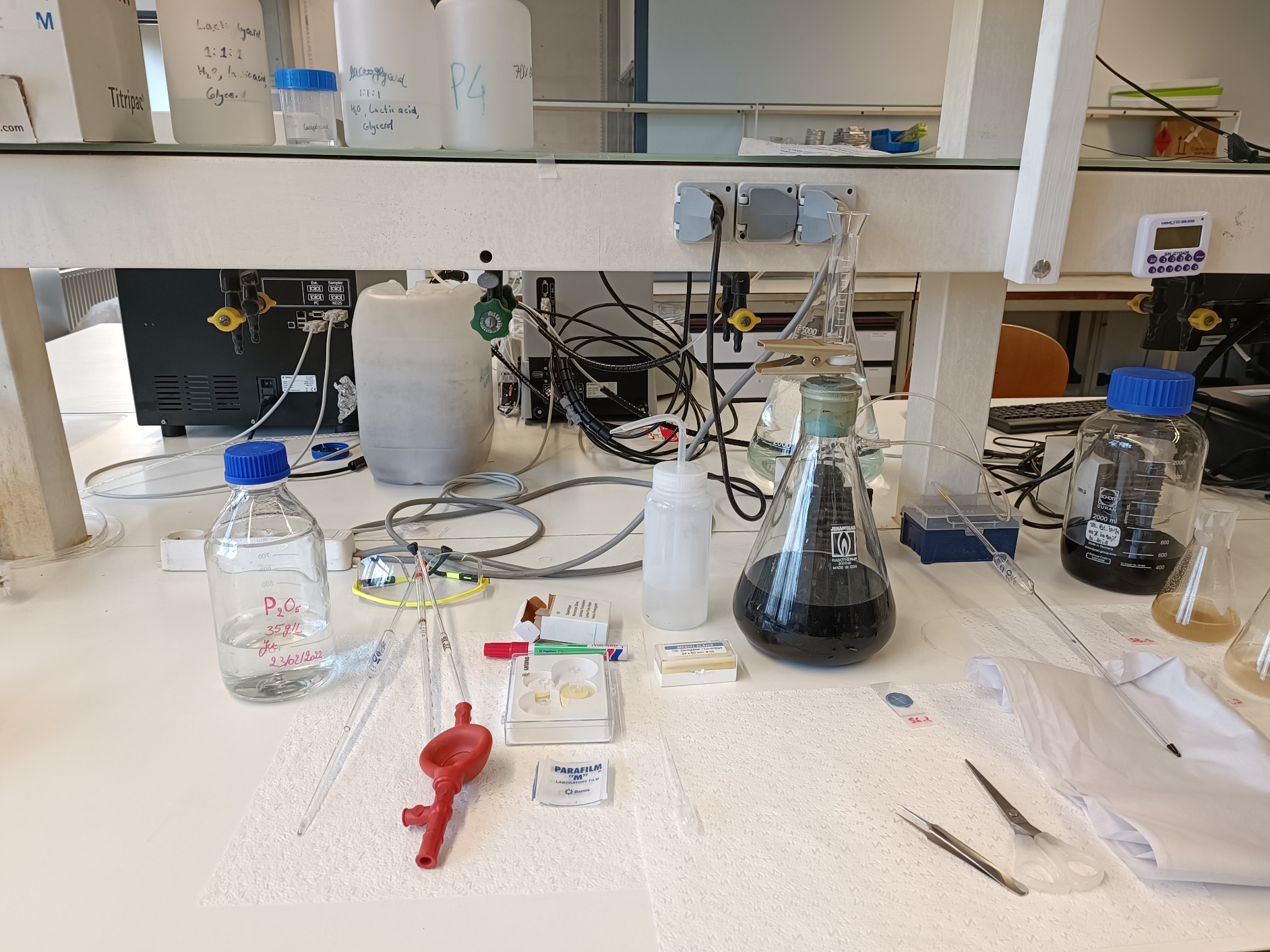

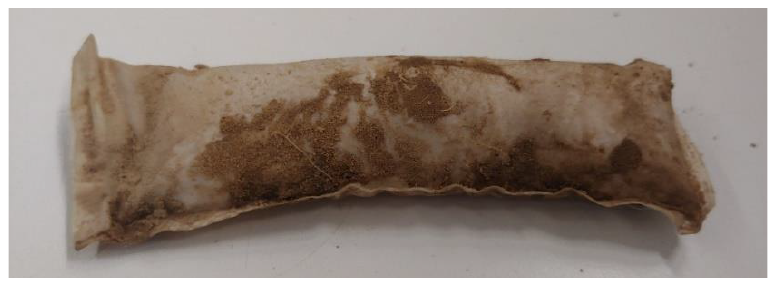

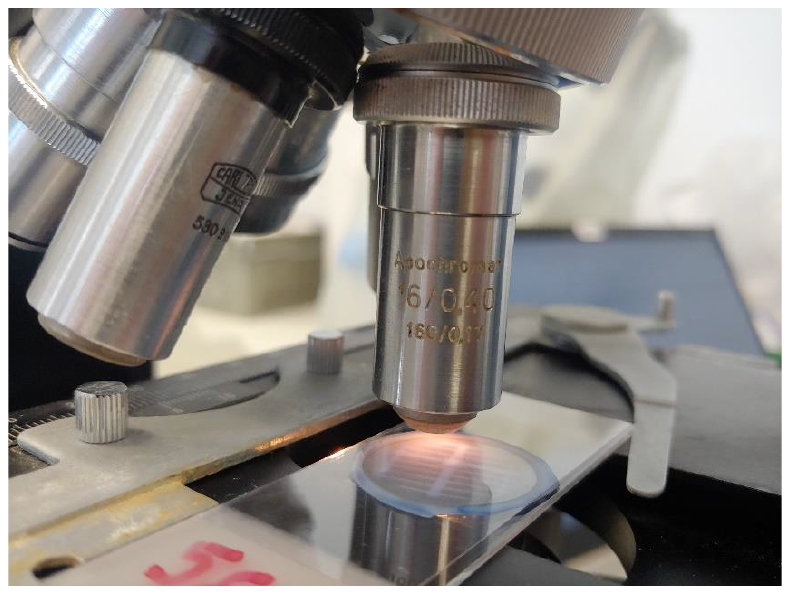

Outlet

Filter

b

a

c

Table S12: Number of mesocosms for which soil heavy metal concentration in each depth and soil fraction combination was below detection limit (bd) or lower than the limit of quantification (LOQ) for each treatment. Number is given compared to the total number of mesocosms of that treatment. LOQ is also given for each element. If none of the samples were bd or <LOQ, a “/” is shown, if all samples were bd or <LOQ, “all” is shown.

|  |  | Al |  | Cr |  | Ni |  | Zn |  |
| --- | --- | --- | --- | --- | --- | --- | --- | --- | --- |
|  |  | bd | <LOQ | bd | <LOQ | bd | <LOQ | bd | <LOQ |
|  |  |  | <187.9 ppm |  | <19.5 ppb |  | <19.4 ppb |  | <37.2 ppm |
| 0-20 cm | Exchangeable | / | / | / | All | B-AMF: 2/5 B+AMF: 0/5 C-AMF: 2/5 C+AMF: 1/5 | B-AMF: 3/5 B+AMF: 5/5 C-AMF: 3/5 C+AMF: 4/5 | / | B-AMF: 0/5 B+AMF: 1/5 C-AMF: 1/5 C+AMF: 0/5 |
|  | Carbonate | / | / | All | All | All | All | / | / |
|  | Oxidisable | / | / | / | B-AMF: 1/5 B+AMF: 2/5 C-AMF: 2/5 C+AMF: 1/5 | B-AMF: 0/5 B+AMF: 0/5 C-AMF: 4/5 C+AMF: 5/5 | B-AMF: 5/5 B+AMF: 5/5 C-AMF: 0/5 C+AMF: 0/5 | All | / |
|  | Reducible | / | / | / | / | / | / | / | / |
| 20-40 cm | Exchangeable | / | / | All | / | B-AMF: 0/5 B+AMF: 0/5 C-AMF: 3/5 C+AMF: 3/5 | B-AMF: 5/5 B+AMF: 5/5 C-AMF: 2/5 C+AMF: 2/5 | / | / |
|  | Carbonate | / | / | All | All | All | All | / | / |
|  | Oxidisable | / | / | B-AMF: 0/5 B+AMF: 0/5 C-AMF: 2/5 C+AMF: 1/5 | / | All | / | All | / |
|  | Reducible | / | / | / | / | / | / | / | / |
| 40-60 cm | Exchangeable | All | / | All | / | All | / | B-AMF: 1/5 B+AMF: 2/5 C-AMF: 1/4 C+AMF: 4/5 | B-AMF: 4/5 B+AMF: 3/5 C-AMF: 3/4 C+AMF: 1/5 |
|  | Carbonate | B-AMF: 1/5 B+AMF: 1/5 C-AMF: 1/4 C+AMF: 1/5 | / | B-AMF: 4/5 B+AMF: 5/5 C-AMF: 4/4 C+AMF: 4/5 | / | B-AMF: 4/5 B+AMF: 5/5 C-AMF: 4/4 C+AMF: 4/5 | / | B-AMF: 1/5 B+AMF: 1/5 C-AMF: 1/4 C+AMF: 1/5 | / |
|  | Oxidisable | / | B-AMF: 5/5 B+AMF: 5/5 C-AMF: 3/4 C+AMF: 4/5 | / | / | / | All | B-AMF: 1/5 B+AMF: 2/5 C-AMF: 1/4 C+AMF: 3/5 | B-AMF: 4/5 B+AMF: 3/5 C-AMF: 2/4 C+AMF: 1/5 |
|  | Reducible | / | / | / | / | / | / | / | / |

Table S13: Concentrations of heavy metals (Al, Cr, Ni, Zn) in the soil at the end of the experiment that were detectable in the layers and fractions for which other measurements were all below the detection limit or lower than the limit of quantification.

| Mesocosm | Fraction | layer | Treatment | element | Concentration  (mg kg^-1^ soil) |
| --- | --- | --- | --- | --- | --- |
| 18 | Carbonate | 40-60cm | CP+A | Cr | 0.96 |
| 18 | Carbonate | 40-60cm | CP+A | Ni | 0.98 |
| 22 | Oxidisable | 40-60cm | CP-A | Al | 45.11 |
| 22 | Oxidisable | 40-60cm | CP-A | Zn | 0.49 |
| 61 | Oxidisable | 40-60cm | CP+A | Al | 45.32 |
| 61 | Oxidisable | 40-60cm | CP+A | Zn | 0.55 |
| 61 | Oxidisable | 0-20cm | CP+A | Ni | 0.53 |
| 62 | Carbonate | 40-60cm | B-AMF | Cr | 0.98 |
| 62 | Carbonate | 40-60cm | B-AMF | Ni | 1.03 |

Table S14: Comparison of average increases (moles charge equivalents mesocosm^-1^) ± SE between the start and the end of the experiment in the soil, and increases in Ca, Mg and K in the plants compared to C-AMF treatment ± SE. As leachates were absent after 20 June 2022, they were not included in weathering rates calculations.

| Treatment | Unit | Soil | Plants |
| --- | --- | --- | --- |
| B+AMF | mol mesocosm^-1^ | 4.12 ± 0.32 | 0.04 ± 0.02 |
| B-AMF | mol mesocosm^-1^ | 3.74 ± 0.35 | 0.02 ± 0.02 |
| C+AMF | mol mesocosm^-1^ | 0.65 ± 0.1 | 0.01 ± 0.02 |
| C-AMF | mol mesocosm^-1^ | 0.67 ± 0.13 | 0 |

Figure S15: Total aboveground dry biomass (left) and aboveground biomass for each organ (right) determined after drying at 70 °C until constant weight for the four treatments (B-AMF = basalt without AMF, B+AMF = basalt with AMF, CP-A = control without AMF, CP+A = control with AMF) at the end of the experiment. Boxes represent the interquartile range (25th–75th percentile), horizontal lines indicate the median, and whiskers extend to the most extreme data points within 1.5× of the interquartile range. Points outside this range are plotted as outliers. p- and F-values from a linear regression analysis with dry tissue biomass as response variables and basalt, AMF and their interaction as covariables are shown. Interactions were not significant and were excluded from the model. Significant relationships are indicated by an asterisk (*).

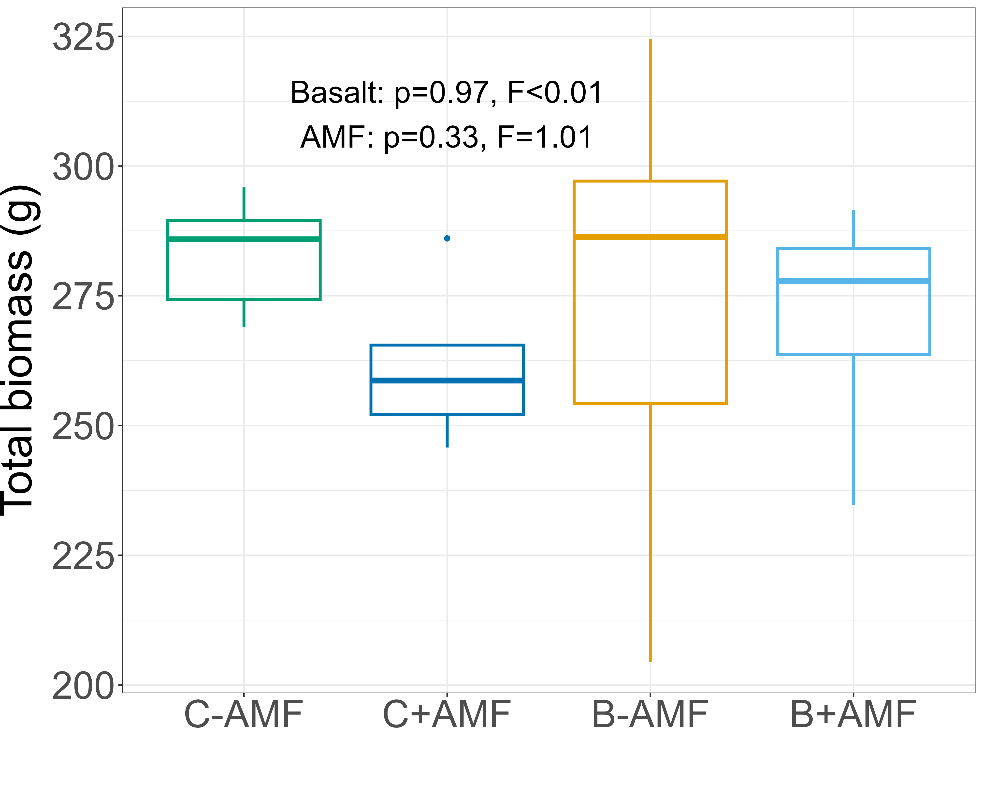

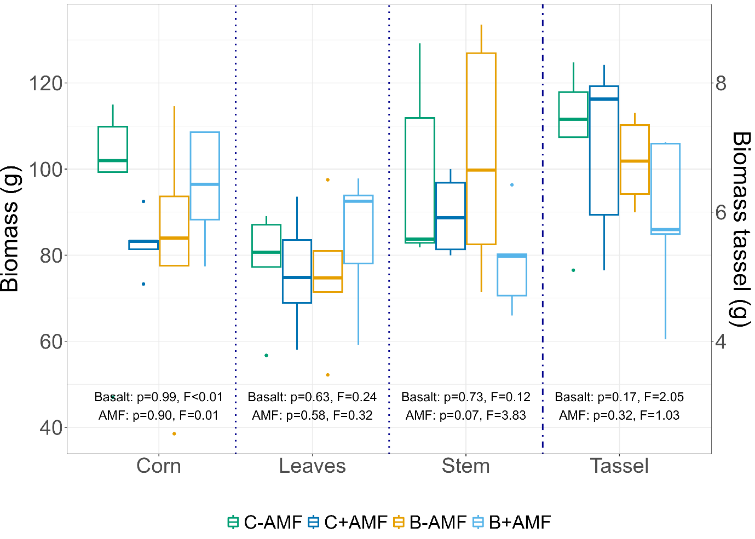

Figure S16: Leaf area index (LAI; left panel) in the middle of the growing season (20 July 2022) for each treatment, determined following Ven et al. (2019). Plant height during the experiment for each treatment (right panel; B-AMF = basalt without AMF, B+AMF = basalt with AMF, CP-A = control without AMF, CP+A = control with AMF). Left panel: p- and F-values are shown from a linear regression analysis with LAI as response variable and basalt, AMF, and their interaction as covariables. The interaction was not significant and was excluded from the model. Significant relationships are indicated by an asterisk (*). Right panel: p- and F-values are shown from a linear regression analysis with plant height as fixed effect and basalt, AMF, time and their interaction as covariables. Interactions were not significant and were excluded from the model.

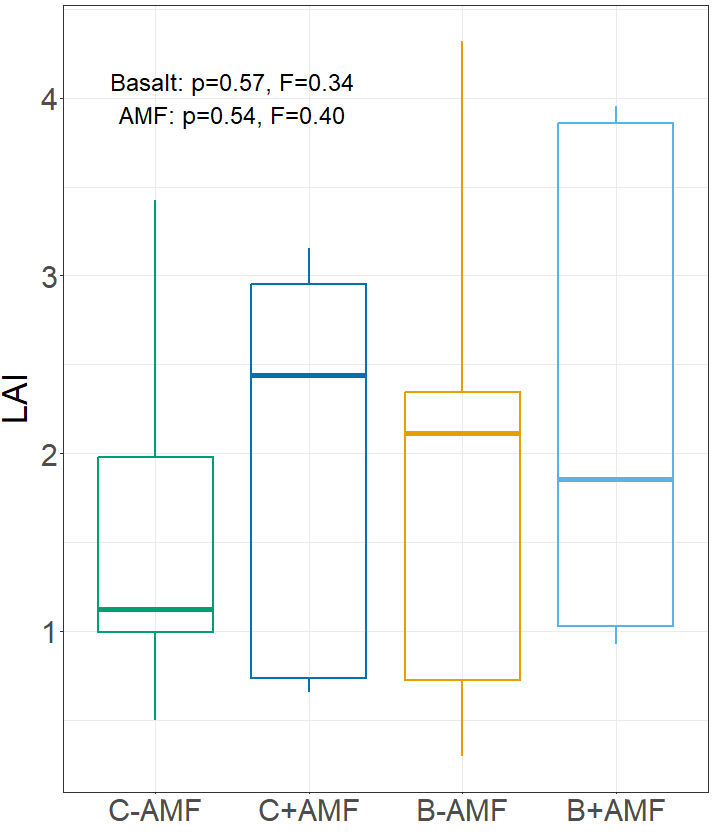

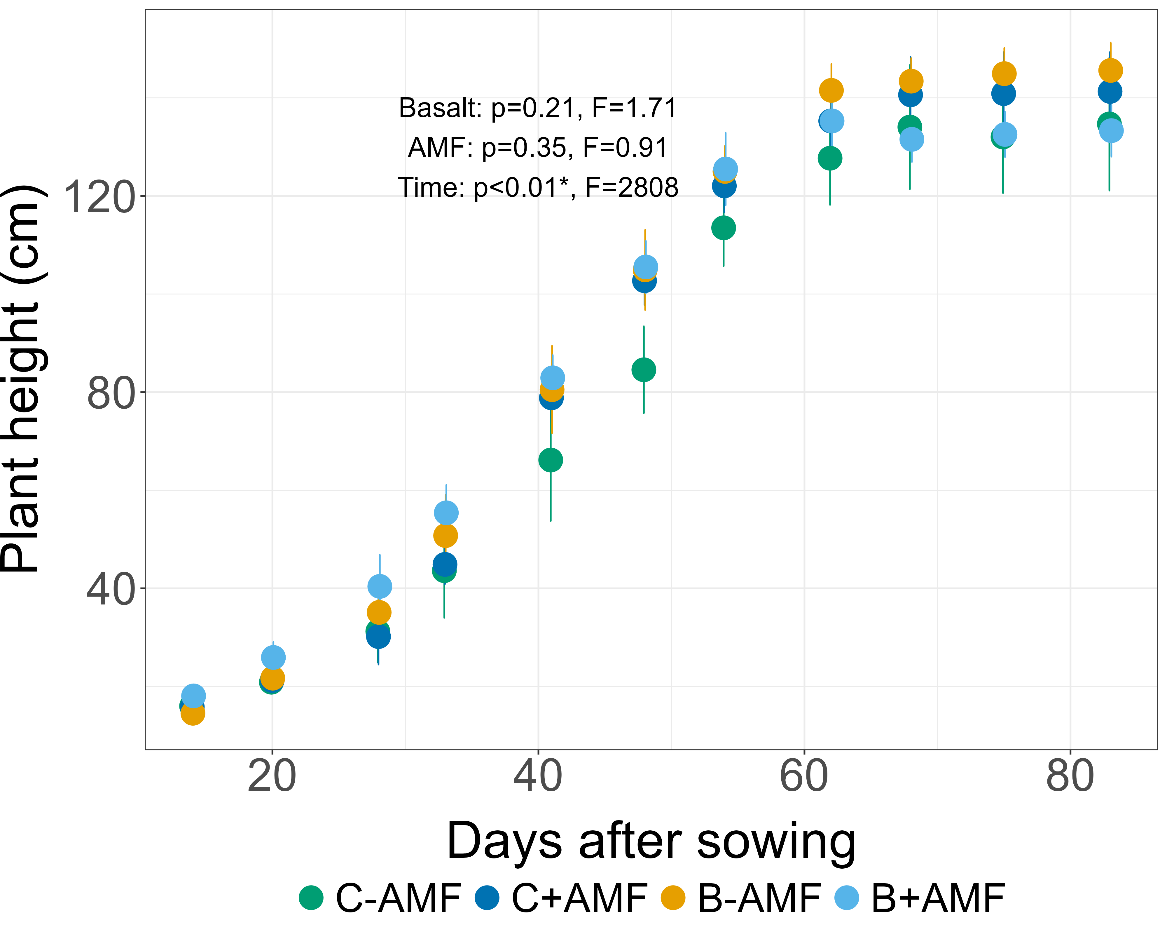

Table S17: p- and F-values from a linear regression analysis with element concentration in various plant parts as response variable, and basalt, AMF and their interaction as covariables. Interactions that were not significant were excluded from the model. Significant relationships are indicated by an asterisk (*). Assumptions of normality or heteroscedasticity were not met for stem and corn C concentrations. In those instances, a non-parametric Kruskal-Wallis test was used instead, with treatment as a covariable. P-and F-value or p- and chi² values are shown. Cd, Cr, and V concentrations in the leaves, and V concentrations in corn were all below the limit of quantification (LOQ). Despite the significant basalt x AMF interaction in stem CN ratio (Fig. S9), the Tukey post-hoc pairwise comparison revealed no significant difference.

|  |  | **Stem** | | **Leaves** | | **Corn** | | **Tassel** | | **Roots** | |
| --- | --- | --- | --- | --- | --- | --- | --- | --- | --- | --- | --- |
|  |  | p-value | F | p-value | F | p-value | F | p-value | F | p-value | F |
| Al | Basalt | 0.62 | 0.25 | 0.23 | 1.58 | 0.23 | 1.56 | 0.69 | 0.16 | 0.62 | 0.25 |
|  | AMF | 0.34 | 0.96 | 0.23 | 1.52 | 0.38 | 0.84 | 0.44 | 0.64 | 0.66 | 0.19 |
| Ca | Basalt | 0.95 | <0.01 | 0.5 | 0.48 | 0.19 | 1.94 | 0.99 | 0 | 0.53 | 0.41 |
|  | AMF | 0.94 | <0.01 | 0.83 | 0.05 | 0.25 | 1.42 | 0.28 | 1.23 | 0.13 | 2.56 |
| Cd | Basalt | 0.42 | 0.69 | <LOQ | | 0.25 | 1.47 | 0.74 | 0.11 | 0.61 | 0.27 |
|  | AMF | **<0.01*** | 10.1 |  |  | 0.42 | 0.7 | 0.47 | 0.55 | 0.66 | 0.20 |
| Cr | Basalt | 0.92 | 0.01 | <LOQ | | 0.35 | 0.93 | 0.19 | 0.86 | 0.49 | 0.5 |
|  | AMF | 0.99 | <0.01 |  |  | **0.03*** | 5.68 | 0.51 | 0.46 | 0.87 | 0.03 |
| Fe | Basalt | 0.34 | 0.97 | 0.21 | 1.69 | 0.65 | 0.22 | 0.41 | 0.7 | 0.49 | 0.5 |
|  | AMF | 0.33 | 0.99 | 0.49 | 0.5 | 0.83 | 0.05 | 0.9 | 0.02 | 0.8 | 0.07 |
| K | Basalt | 0.27 | 1.32 | **0.03*** | 5.51 | 0.23 | 1.54 | 0.84 | 0.04 | 0.96 | 0.03 |
|  | AMF | 0.38 | 0.81 | 0.74 | 0.12 | 0.41 | 0.70 | 0.81 | 0.06 | 0.62 | 0.26 |
| Mg | Basalt | **<0.01*** | 14.7 | **<0.001*** | 19.02 | 0.26 | 1.39 | **<0.01*** | 12.59 | **<0.001*** | 36.3 |
|  | AMF | 0.88 | 0.02 | 0.32 | 1.07 | 0.47 | 0.55 | 0.06 | 4.18 | **<0.01*** | 11.6 |
|  | B^AMF | ns | ns | ns | ns | ns | ns | ns | ns | *0.08* | 3.47 |
| Ni | Basalt | 0.58 | 0.31 | 0.1 | 3.01 | **0.02*** | 7.21 | 0.09 | 3.19 | 0.69 | 0.17 |
|  | AMF | 0.99 | 0 | 0.97 | <0.01 | **0.03*** | 5.63 | 0.37 | 0.86 | 0.94 | <0.01 |
| P | Basalt | 0.89 | 0.02 | 0.75 | 0.11 | 0.90 | 0.2 | 0.37 | 0.84 | 0.34 | 0.94 |
|  | AMF | 0.20 | 1.79 | **0.04*** | 5.2 | 0.21 | 1.74 | 0.38 | 0.83 | 0.21 | 1.73 |
| Pb | Basalt | 0.11 | 2.87 | **0.03*** | 5.36 | 0.67 | 0.19 | 0.37 | 0.86 | **<0.01*** | 14.9 |
|  | AMF | 0.52 | 0.43 | 0.68 | 0.17 | 0.62 | 0.26 | 0.91 | 0.01 | 0.48 | 0.52 |
|  | B^AMF | ns | ns | ns | ns | ns | ns | ns | ns | **0.048*** | 4.63 |
| V | Basalt | 0.95 | <0.01 | <LOQ | | <LOQ | | 0.59 | 0.3 | 0.35 | 0.93 |
|  | AMF | 0.24 | 1.5 |  |  |  |  | 0.86 | 0.03 | *0.09* | 3.19 |
|  | B^AMF | ns | ns |  |  |  |  | ns | ns | *0.054* | 4.36 |
| Zn | Basalt | 0.75 | 0.1 | 0.62 | 0.25 | 0.15 | 2.27 | 0.75 | 0.1 | 0.36 | 0.89 |
|  | AMF | 0.53 | 0.4 | 0.58 | 0.31 | 0.63 | 0.25 | 0.21 | 1.74 | 0.28 | 1.24 |
| C | Basalt | Kruskal-Wallis test: | | **0.02*** | 6.87 | Kruskal-Wallis test: | | Kruskal-Wallis test: | | 0.46 | 0.57 |
|  | AMF | P=0.50 | chi²= 2.35 | 0.19 | 1.85 | p=0.87, chi²=0.73 | | p=0.31, chi²=3.57 | | 0.66 | 0.20 |
| N | Basalt | 0.57 | 0.33 | 0.16 | 2.15 | 0.86 | 0.03 | 0.84 | 0.04 | 0.61 | 0.27 |
|  | AMF | 0.52 | 0.44 | 0.12 | 2.76 | 0.62 | 0.26 | *0.05* | 4.34 | *0.09* | 3.16 |
| Si | Basalt | **<0.01*** | 9.08 | 0.19 | 1.83 | 0.14 | 2.41 | **0.02*** | 6.9 | 0.34 | 0.98 |
|  | AMF | 0.63 | 0.24 | 0.60 | 0.28 | 0.88 | 0.03 | 0.47 | 0.55 | 0.41 | 0.72 |
| CN ratio | Basalt | 0.45 | 0.61 | 0.17 | 2.01 | 0.9 | 0.02 | 0.98 | 0.0006 | 0.88 | 0.03 |
|  | AMF | 0.5 | 0.47 | 0.22 | 1.60 | 0.55 | 0.37 | **0.02*** | 6.21 | *0.09* | 3.23 |
|  | B^AMF | **0.047*** | 4.62 | ns | ns | ns | ns | ns | ns | ns | ns |

Figure S18: Mean dissolved organic carbon (DOC) in the porewater over the experimental period in the four treatments which used pasteurized soil (C-AMF, C+AMF, B-AMF, B+AMF) compared to a parallel mesocosm experiment at the same location which used the same basalt and soil, without pasteurization. Pasteurized treatments are indicated by a triangle, unpasteurized by a circle. Error bars represent standard errors on the mean.

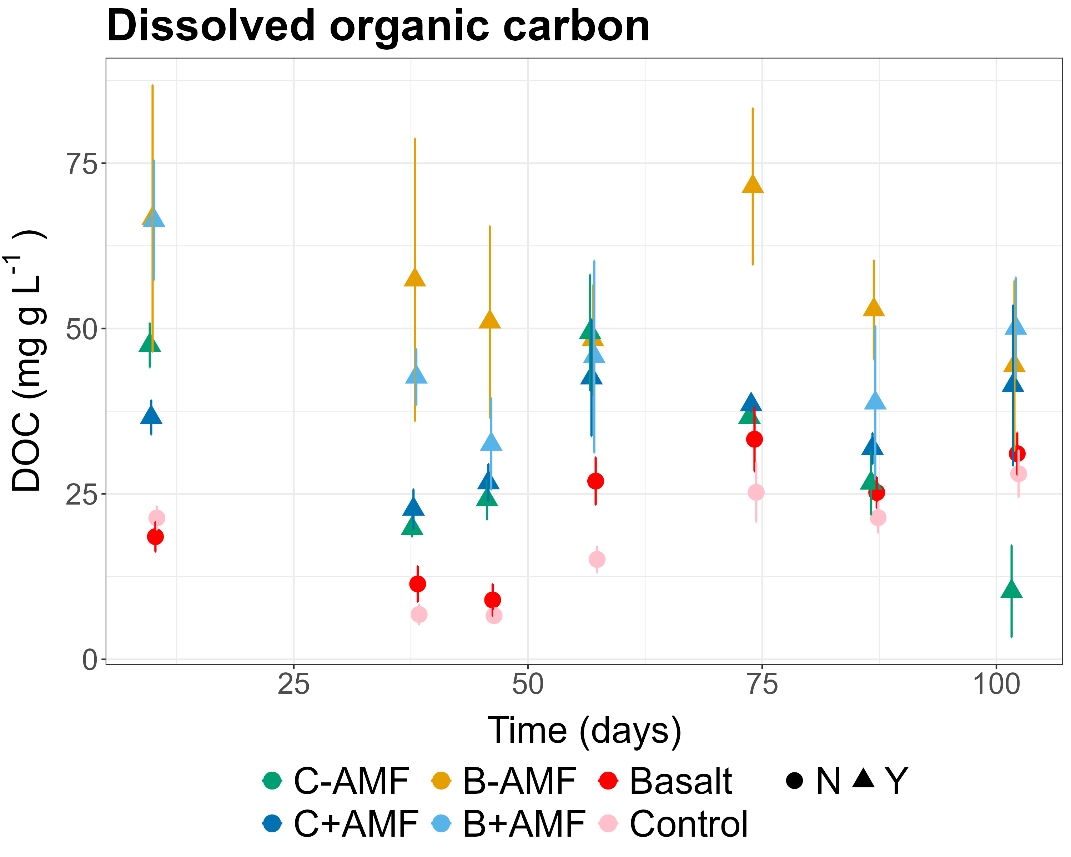

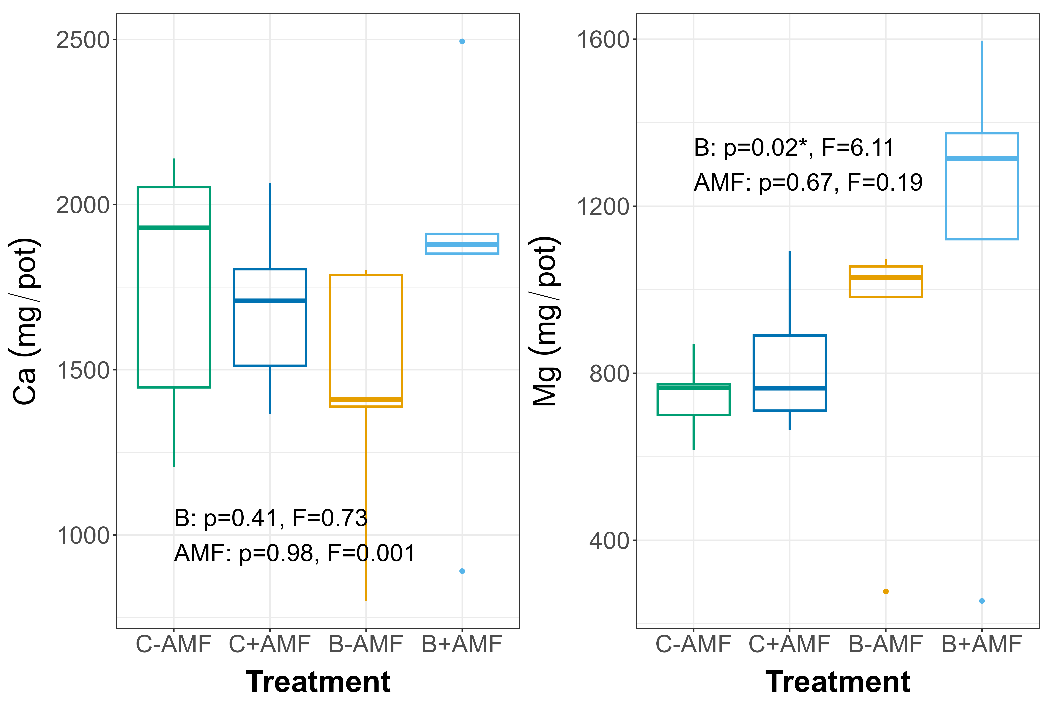

Figure S19: Ca and Mg stocks in the plant for each treatment (C-AMF, C+AMF, B-AMF, B+AMF) calculated by multiplying each cation concentration in each plant tissue (corn, stem, root, leaves, tassel) by the biomass of said plant tissue and then summating the stocks in each plant tissue. Boxes represent the interquartile range (25th–75th percentile), horizontal lines indicate the median, and whiskers extend to the most extreme data points within 1.5× of the interquartile range. Points outside this range are plotted as outliers. p- and F-values are shown from a linear regression analysis with stock as response variable and basalt, AMF, and their interaction as covariables. The interaction was not significant and was excluded from the model. Significant relationships are indicated by an asterisk (*).

Figure S20: soil exchangeable bases (Ca, K, Mg and Na) for each treatment (B-AMF = basalt without AMF, B+AMF = basalt with AMF, CP-A = control without AMF, CP+A = control with AMF). p- and F-values are shown from a linear regression analysis with the exchangeable bases as response variables and basalt, AMF and their interaction as covariables. Interactions were not significant and were excluded from the model. Significant relationships are indicated by an asterisk (*).

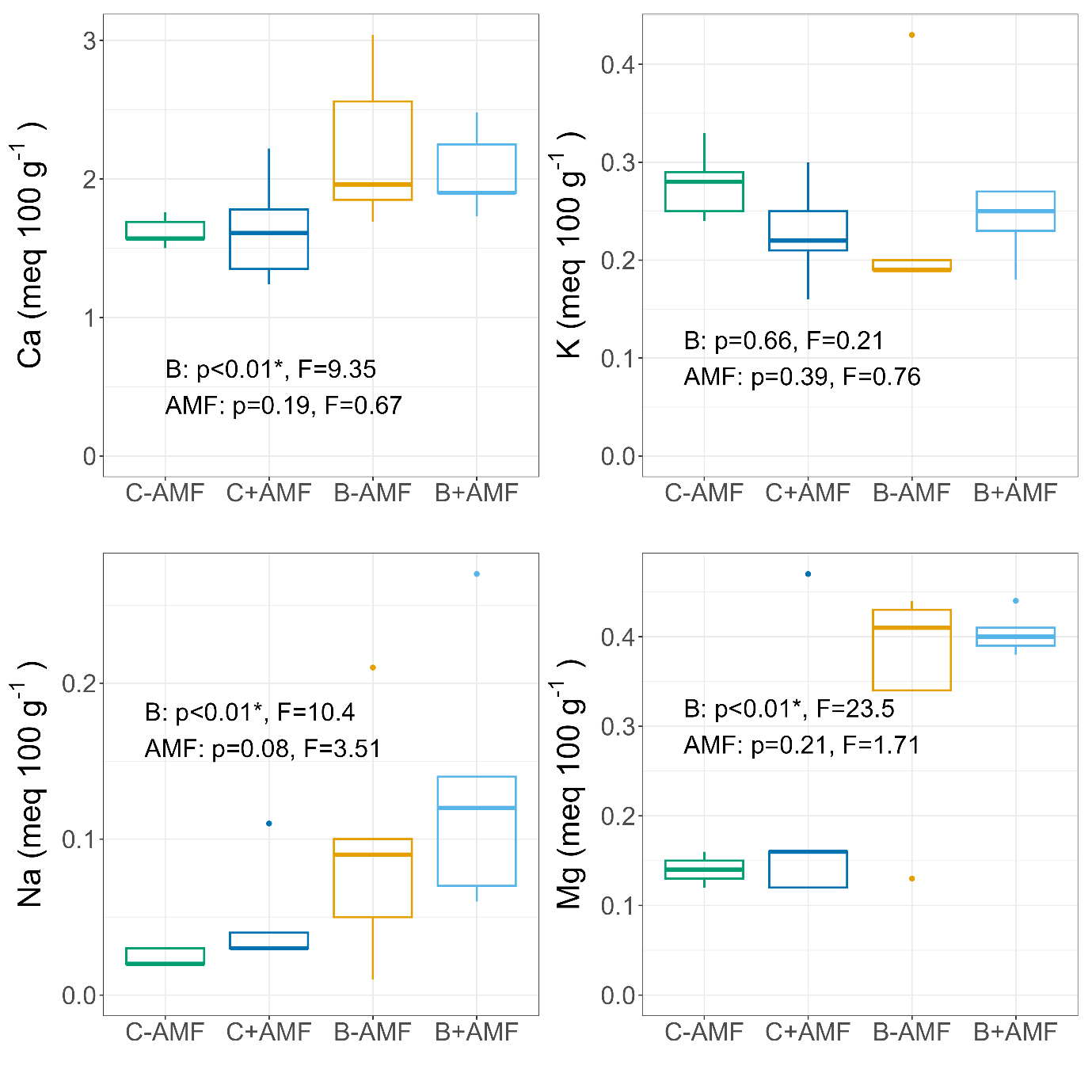

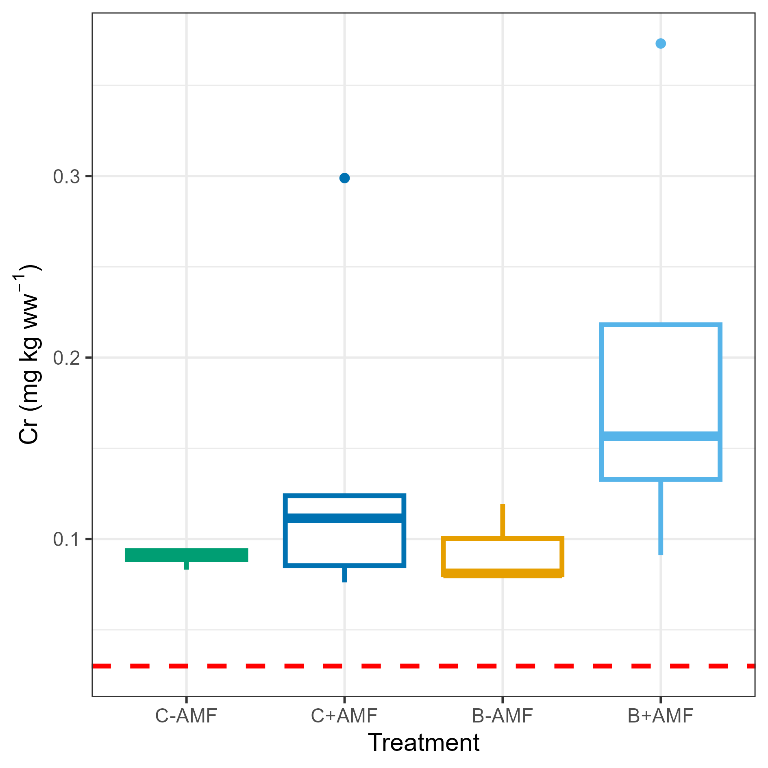

Figure S21: Concentration of Cr in the corn of *Zea mays* for each treatment (C-AMF = control without AMF, C+AMF = control with AMF, B-AMF = basalt without AMF, B+AMF = basalt with AMF) compared to WHO guidelines (dashed red line). Boxes represent the interquartile range (25th–75th percentile), horizontal lines indicate the median, and whiskers extend to the most extreme data points within 1.5× of the interquartile range. Points outside this range are plotted as outliers.
